# Behavioral read out of population value signals in primate orbitofrontal cortex

**DOI:** 10.1101/2021.03.11.434452

**Authors:** Vincent B. McGinty, Shira M. Lupkin

## Abstract

The primate orbitofrontal cortex (OFC) has long been recognized for its role in value-based decisions; however, the exact mechanism linking OFC value representations to decision outcomes has remained elusive. To address this question we show for the first time that trial-wise variability in choices can be explained by variability in value signals decoded from many simultaneously recorded OFC neurons. Mechanistically, this relationship is consistent with the projection of activity within a low-dimensional value-encoding subspace onto a potentially higher-dimensional, behaviorally-potent output subspace.

## INTRODUCTION

One of the most powerful ways of identifying decision mechanisms is to show a relationship between neural variability and behavioral variability in single trials. A canonical example is in the primate extrastriate area MT, where trial-to-trial variability in the firing of motion-sensitive neurons correlates with variability in motion judgements, even over trials in which the same noisy motion stimulus is repeated many times^1^. For any form of decision behavior, identifying such a fine-grained neural-behavioral link is a *sine qua non* for understanding its neural mechanism; it is not only a necessary (though not sufficient^2–4)^ condition for demonstrating a causal role of neural activity for behavior, but it also permits computational insights into how activity in a given area is interpreted by downstream circuits and ultimately transformed into action.^5–8^

For economic choices, defined as those based upon value or preference, the OFC is thought to have a critical causal role: encoding evidence in the form of value representations that are ultimately transformed into a decision^9–13^. Substantial converging evidence supports this hypothesis. First, perturbation of OFC activity disrupts value-based choices^14–19^. Second, a large fraction of OFC neurons encodes economic value^9–11, 20, 21^, a function for which this region appears to be specialized within the frontal lobe^22^. Third, the sensitivity of value coding neurons to different offers matches the behavioral sensitivity of the subjects measured in the same sessions^11^, and that network-level fluctuations in OFC value representations correlate with decision reaction times (though not with the choices themselves) in single trials^12^. However, while these studies suggest some link between value representation and decision behavior, one key finding has been conspicuously absent: Even in large and rigorously analyzed samples of OFC cells, a significant trial-to-trial correlation between value coding in single neurons and value-based choices has yet to be found^23–25^. This is especially puzzling; if OFC value signals mediate value-based choices, there must be some measurable relationship between them. Moreover, it is a major barrier to progress in the field, because until this relationship is identified, it will be difficult to fully validate the causal role of OFC, or to identify the mechanisms by which value signals are read out downstream and transformed into action – such as mechanisms already identified for perceptual choices.

One potential explanation, suggested by the elegant work of Conen and Padoa-Schioppa^25^, is that the choice-predictive activity is difficult to detect in single OFC cells because of the low degree of shared noise in the population (for more see also Haefner et al.^26^). We reasoned that if this hypothesis were true, more informative choice-predictive signals may be obtained by pooling the value representations of many simultaneously recorded neurons in single trials. Such multivariate signals inherently convey more information than univariate signals, provided that the individual contributors to the signal (single neurons in our case) are sufficiently independent^12, 27, 28^. To test this hypothesis, we recorded from populations of OFC neurons in monkeys making value-based choices. Consistent with prior studies, we identify population-level representations of economic value^12, 22, 24, 29^; however, unlike prior work, we use the pooled activity of many cells to decode single-trial estimates of the offer values, and identify a robust relationship between these value representations and the decisions made in individual trials. In addition, we leverage the population nature of the approach to identify for the first time a behavioral output subspace within OFC (i.e. the multivariate patterns of neural activity that maximally explain variance in choice) and characterize its key properties, as well as its relationship with the neural subspace encoding offer values.

## RESULTS

Two macaque monkeys performed a novel economic choice task. In every trial, two choice targets were shown on the left and right sides of the task display, each associated with the delivery of a fixed juice reward, between 1-5 drops (Fig. 1a,b). There were 12 physically unique targets used in a given session, but only 5 levels of juice reward; the term ‘target identity’ therefore refers to a target’s unique physical appearance, and ‘offer value’ or ‘target value’ is used to refer to its reward association. An important feature of this task is that after an initial fixation period, saccadic eye movements were unrestricted – and in fact were required for optimal performance. The reason is that in each trial the two choice targets were obscured from view until the monkeys fixated upon them directly, an effect achieved through the use of visual crowding (Fig. 1c)^30, 31^ and a dynamic display that was contingent upon gaze (see Methods, and also Hunt et al.^22^ for a similar approach). The monkeys therefore viewed the targets sequentially, by fixating them one at a time. Monkeys could view the targets in any order and for any amount of time, and eye tracking was used to determine when each target was viewed. The monkeys were free to choose at any time, and did so by first lifting a central response lever (extinguishing the targets), and then pressing the left or right lever within a 500ms deadline (Fig. 1a, bottom). The task therefore combines two features that were advantageous for neural analysis: First, in each trial the decision time gives an upper temporal boundary for pre-decision neural activity. Second, the target viewing times indicate when the offer values become known to the monkey in each trial; all neural analyses were therefore performed on data that were time-locked to the two target viewing times in each trial – e.g. see x-axis labels in Figs. 2d-e, 3a-b, and 6c.

**Figure 1:**
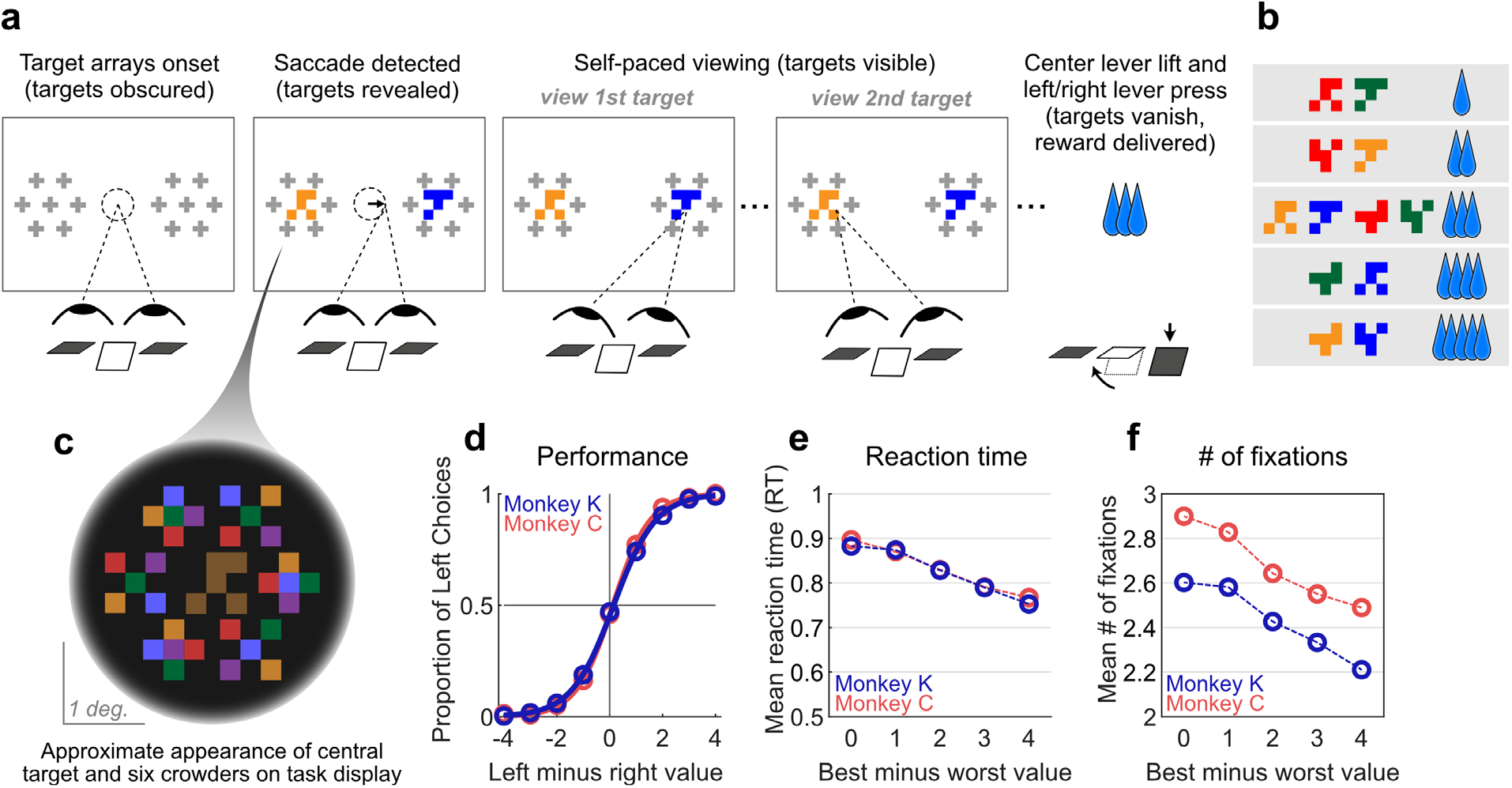
Decision-making task and performance in two monkeys. **(a)** Abbreviated task sequence, not shown to scale; see Extended Data Fig. 1 for full task sequence. The yellow and blue glyphs are choice targets, and the six gray “+” shapes indicate the location of crowders designed to obscure the targets until they are viewed (fixated) directly. For clarity, in this panel the crowders are illustrated in gray at reduced scale; on the actual task display, crowders were multicolored and the same size as the targets, as in panel (c). The interval between target arrays onset and center lever lift defines the decision reaction time (RT). **(b)** Example set of 12 choice targets, organized into 5 groups corresponding to the 5 levels of associated juice reward. (**c**) Close-up view of a single target array, consisting of a central yellow choice target surrounded by six non-task-relevant crowders. Two such arrays appear on the display, each centered 7.5° from the display center. **(d-f)** Choices, reaction time, and fixations over 10,972 trials in 16 sessions for Monkey C and 9,617 trials in 16 sessions for Monkey K. **(d)** Fraction of left choices as a function of the left minus right offer value. Smooth lines give logistic fit. Logistic regression estimates for effect of left minus right value: Monkey K, 1.25 (SE 0.02, n = 9,617 trials); Monkey C, 1.42 (SE 0.03, n = 10,972 trials). **(e)** Mean reaction times decreased as a function of trial difficulty, defined as the absolute difference in offer values. SEMs are smaller than 0.004s for all data points. A linear fit of RT as a function of difficulty gives the following average slopes: Monkey K −0.035 (SE 0.001); Monkey C, −0.035 (SE 0.001). **(f)** Mean number of fixations performed per trial, as a function of trial difficulty. SEMs are smaller than 0.03 for all data points. A linear fit gives the following average slopes: Monkey K, −0.106 (SE 0.007); Monkey C, −0.120 (SE 0.006).

**Figure 2:**
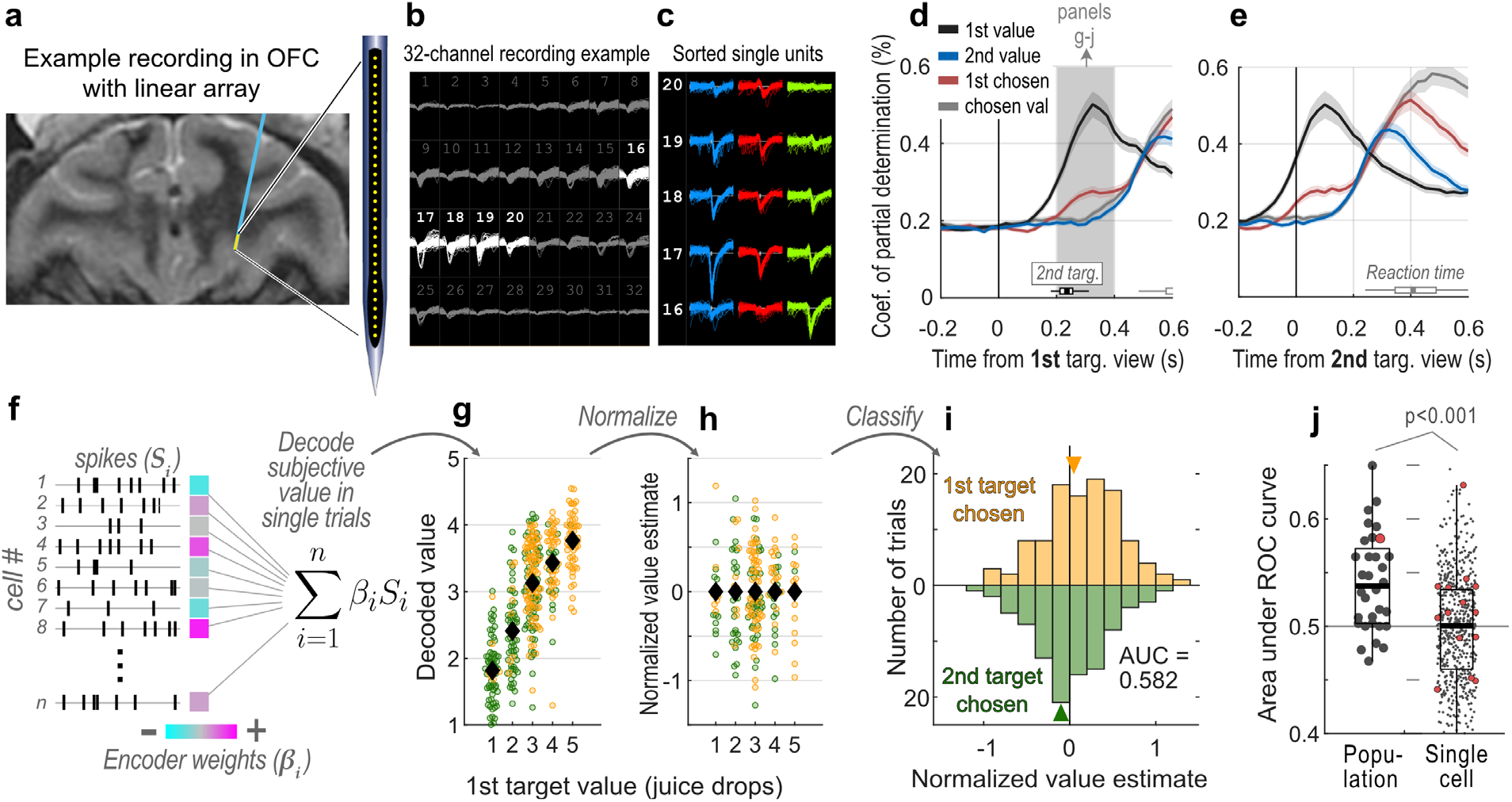
OFC recording and population decoding of choice. **(a)** Example placement of electrode array in OFC. See also Extended Data Fig. 2. **(b)** Example traces. Numerals give array channels. Highlighted channels are shown in panel (c). **(c)** Example of three sorted units. **(d-e)** Encoding of task variables in single cells, measured by coefficients of partial determination in a linear model (Eqn. 1). In (d) and (e), the data are aligned to the viewing of the first and second target in each trial, respectively. Lines and shaded area give the mean and SEM over 1450 cells. The black and gray box plots above the x-axis indicate the onsets of second target viewing and of the decision RT, respectively. The shaded interval gives the epoch used in panels (g-j). **(f)** Schematic of choice decoding based on population-level representation of the first offer value. Cyan-magenta scale illustrates cell weights obtained from a linear model to explain the first target value over training trials (Eqn. 2). (**g**) Decoded estimates of the value of the first target over test trials from a single session. Each dot gives weighted sum of spiking observed across all recorded cells in one test trial (Eqn. 3, n = 284 test trials total). Colors indicate trials in which the first or second target was chosen (yellow and green, respectively). Trials are grouped according to the first offer value; black diamonds indicate group means, and horizontal spread within groups is for visualization only. **(h)** Normalized value estimates (NVEs), obtained by residualizing the decoded values in panel (g) with respect to the target identities. NVEs quantify the degree to which the decoded values are more or less than expected given the target identity. Only test trials with value differences of 0 or 1 are shown (n = 141 out of 284 trials). Conventions are the same as in panel (g). **(i)** Variability in the NVEs predicts variability in choices. Data in panel (h) were divided according to the decision outcome (first/second target chosen in the upper/lower histograms, respectively). Arrows give the distribution medians. **(j)** Accuracy of choice-predictive activity aggregated across sessions and monkeys. On average, choice probabilities derived from population-based NVEs (left, 32 sessions) were significantly larger than choice probabilities derived from single cells (right, 599 cells with non-zero weights in Eqn. 2). Red dots indicate data from session shown in panels (g-i). Gray line indicates chance-level performance. P-value reflects two sample t-test, with p = 4.6e-6, t_629_ = 4.62.

**Figure 3:**
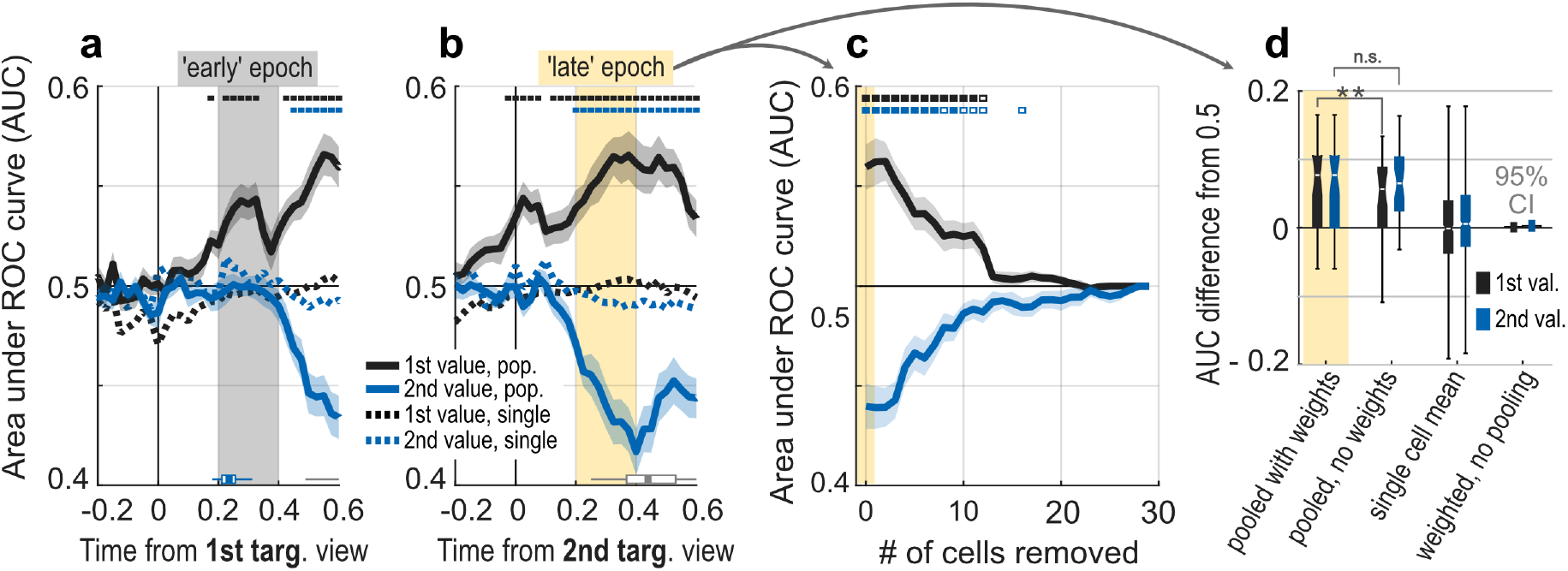
Decoding choice over time, neuron dropping, and controls. **(a-b)** Data are time-locked to target viewing times. Population-based choice decoding is given by solid lines with shading (means and SEMs, 32 sessions). Small rectangles at top indicate bins for which the population effects are significantly different from 0.5 at a corrected threshold of p<0.05 using a one-sample t-test. Dotted lines give the means of choice probabilities calculated from single cells individually (no SEM shown, numbers of cells vary by bin, see Methods). The blue and gray horizontal box plots give the distribution of second target viewing times and choice RTs, respectively. The shaded epochs in panels (a) and (b) indicate epochs used in subsequent analyses (see main text). **(c)** Neuron dropping analysis using data from late epoch. Colors indicate value variables. Lines and shading give mean and SEM over 32 sessions. Open and closed rectangles indicate significant difference from 0.5 at uncorrected thresholds of p < 0.05 and p < 0.005 (respectively) by t-test. See Extended Data Fig. 5a-h for data from individual monkeys. **(d)** Control analyses using data from the late epoch. Colors indicate value variables. “Pooled with weights” gives the distribution from the late epoch in panel (b) (n=32 sessions); “pooled, no weights” gives results from a decoder using signed weights only (n=32); “single cell” gives the unweighted choice probabilities (n= 531 (black) and 551 cells (blue)). “Weighted, no pooling” gives the 95% CI of the weighted average of the single-cell choice probabilities (bootstrapped 95% CIs for 531 (black) and 551 cells (blue)). Boxplot whiskers give the full data range. ** indicates p = 0.003, t_31_ = 3.24, and n.s. indicates p = 0.64, t_31_ =-0.48 by paired t-test.

The monkeys viewed both targets at least once in the vast majority of trials (99.3% for Monkey C, 93.2% for Monkey K), and their choices were consistent with both offer values being considered in nearly every trial. Their choices were nearly optimal when the offers differed by 3 or 4 drops of juice, but were highly variable when the offers were equal in value (Fig. 1d). In addition, the decision reaction times (RTs) and the mean number of on-target fixations per trial both decreased as a function of the difference in offer values (Fig. 1e,f). The graded nature of the choice outcomes, RTs, and fixations are consistent with a deliberative process dependent upon the offer values^32^. Moreover, because the stimuli were designed to be perceptually uniform, the extremely low rate of suboptimal choices in easy trials indicates that the monkeys did not have difficulty perceiving the stimuli.

OFC neural activity was recorded using multi-channel linear arrays (Fig. 2a-c, Extended Data Fig. 2), and cells were isolated using semi-automated spike-sorting procedures (see Methods). On average, 45 cells were isolated per session (16 sessions and 848 cells for Monkey K, 16 sessions and 602 cells for Monkey C).

Because the offer values in each trial were revealed serially by sequential viewing (Fig. 1a), all neural analyses were performed on data referenced to the target viewing times, and used variables defined in a viewing-order-based reference frame, consistent with prior studies using serial designs^22, 33, 34^. The offer values were characterized by the juice rewards associated with the first and second target viewed by the monkey in each trial (variable names: ‘1st value’ and ‘2nd value’), in units of juice drops (Fig. 1b). Choice outcome was characterized by the variable ‘1st chosen’, (*true* when the first-viewed target was chosen, *false* otherwise), and ‘chosen value’, defined as the value of the target chosen in each trial. So that all variables were defined for every trial, neural analyses used only trials in which monkeys viewed both targets at least once (see above).

The single-cell representation of these variables was quantified using the coefficient of partial determination (CPD), which measures for every cell the unique contribution that each variable makes towards explaining neural activity over and above the contributions of the other variables^22^. Consistent with prior work^22^, the mean CPD for ‘1st value’ increased following the viewing of the first target (black line, Fig. 2d); and for ‘2nd value’ increased following the viewing of the second target (blue line, Fig. 2e). Notably, after viewing the second target, significant representation of ‘1st value’ persisted, (black line in Fig. 2e), consistent with prior findings ^22, 33, 34^. Representation of ‘chosen value’ increased after the viewing the second target (gray line, Fig. 2e), and reached a peak shortly after the median decision reaction time (gray boxplot). The CPD for the variable ‘1st chosen’ initially increased above pre-trial baseline after the first target was viewed, and then increased again following second target viewing (red line, Fig. 2d,e). Although nontrivial encoding of values and choices could also be measured using variables in a spatial-based reference frame, these variables explained less single-cell variance than order-based variables (Extended Data Fig. 3).

Prior studies have shown only a negligible association between the activity of single-value coding neurons and choice outcomes in individual trials^23, 25^. We reasoned that more robust choice-predictive signals may be obtained by pooling the activity of the many simultaneously recorded neurons, given that pooled neural signals are inherently more informative than single neurons, provided that their shared noise is low^8, 27, 28^. We first considered neural activity within the shaded epoch in Fig. 2d, in which ‘1st value’ is strongly represented. To find a population-level (pooled) representation of ‘1st value’ in this epoch, we first used a set of training trials to fit a linear model that explains ‘1st value’ as a function of the activity of all the cells recorded in a given session (Eqn. 2). The fitted coefficients are taken as weights that quantify each cell’s unique contribution to explaining variance in ‘1st value’. Note that L1-regularization (LASSO)^35^ was used, so that cells contributing negligibly to model fit were assigned weights of exactly zero. Then, in a set of held out test trials, the firing of each cell was multiplied by its respective weight and summed (Eqn. 3, Fig. 2f).

The weighted sums provide for every trial a neurally-derived estimate of the value of the first offer. Naturally, as the reward association of the first target increases, the neurally-decoded estimate of its value also increases, evident in the means of these estimates when they are grouped by ‘1st value’ (Fig. 2g). These neurally decoded estimates can be used to classify the offer values in held out trials, with an average area under the ROC curve of ∼0.6 for classification of adjacent value levels (e.g. 2 drops vs. 3 drops, see Extended Data Fig. 4).

Note that over repeated presentations of targets with the same value, the neurally-decoded estimates have substantial across-trial variability, evident in the wide vertical spread *within* each of the groups in Fig. 2g. If this across-trial variability reflects fluctuations in the subjective value that the monkeys assign to the first offer in each trial, then it should explain (i.e. predict) trial-wise variability in the monkeys’ choices. To test this hypothesis, the weighted sums from Eqn. 3 were transformed into *normalized value estimates* (NVE). NVEs measure the degree to which the decoded values exceed or fall short of their expected magnitude given the identity of the target (see Methods). See Fig. 2g and 2h for examples of raw decoded values and corresponding NVEs from a single session.

The NVEs were then used to classify trials according to whether the first or second offer was chosen, using a receiver operating characteristic analysis (ROC). This is illustrated in Fig. 2i: in this example session higher NVEs for ‘1st value’ predicted a greater likelihood of choosing the first target, and lower NVEs predicted a greater likelihood of choosing the second target, corresponding to an area under the ROC curve of 0.582. Importantly, only difficult trials were used for classification – those in which the targets differed in value by 0 or 1 – because these were the only trials in which choices were sufficiently variable (Fig. 1d).

On average the mean area under the ROC curve for the population-derived value signal from the shaded epoch in Fig. 2d was 0.541 (SEM 0.008) significantly greater than chance (p = 1e-4, t_31_ = 5.11, n=32 sessions, Fig. 2j.). Note that these predictive effects occur well before the decision reaction time, and before the value of the other offer is represented in OFC. It is therefore unlikely that they reflect a commitment to the decision; rather, this predictive signal is consistent with the representation of the subjective value of the offer as it fluctuates from trial-to-trial.

Choice decoding was performed over the entire time course of the trial in 200ms bins with 25ms increments, using models fit separately to the variables ‘1st value’ and ‘2nd value’. As shown in Fig. 3a,b, significant choice decoding from the two models was evident starting ∼200ms after viewing the respective targets, mirroring encoding these variables by single cells (Fig. 2d,e). The opposite signs of the ‘1st value’ and ‘2nd value’ effects indicate that higher ‘1st value’ NVEs predict more choices in favor of the first target, whereas higher ‘2nd value’ NVEs predict the opposite. The results were similar when each monkeys’ data was analyzed individually; when using non-condition-normalized data to calculate AUCs^36^; when using data referenced to the decision RT; and when decoding only trials with equal-value offers (Extended Data Fig. 5). Results were also similar when using Ridge or ordinary least squares for fitting and decoding (not shown).

Two follow-up analyses indicate that the predictions made by ‘1st value’ and ‘2nd value’ models were independent. These analyses used data within the ‘late’ epoch indicated in Fig. 3b (200-400ms after viewing the second offer), when decoding for both models was near maximum. First, for every session logistic regression was used to explain choice outcome (first or second offer chosen) as a function of the late epoch NVEs for ‘1st value’ and ‘2nd value’; the model also includes the difference in offer values and nuisance regressors accounting for choice history and other variables (Eqn. 4). On average over 32 sessions, the regression estimates for both the ‘1st value’ and ‘2nd value’ NVEs were significantly different from zero, and were nearly equal magnitude but of opposite signs (mean estimate for ‘1st value’ was 0.240 SEM 0.06, p = 0.0004, t_31_=4.; mean for ‘2nd value’ was −0.234 IQR 0.062, p = 0.0007, t_31_ = −3.78.). This indicates that both neural value signals explain variability in choice when included in the same neural/behavioral model. Estimates were no different in a model without nuisance regressors, and were equal in magnitude on trials in which the offers differed by 1 drop vs. trials in which they were equal in value (Extended Data Fig. 6).

Second, the cell weights from the ‘1st value’ model (Eqn. 2) were uncorrelated with the weights from the ‘2nd value’ model measured over the same epoch (Fig. 4a-b). Likewise, the late epoch NVEs from the two models showed only a small negative correlation (Fig. 4c-d). This indicates that ‘1st value’ and ‘2nd value’ were represented independently, both in terms of the neural activity patterns that represent the values, and in terms of the trial-wise fluctuations of the value signals (the NVEs). Taken together, these results show that the OFC maintains highly separable representations of ‘1st value’ and ‘2nd value’, and that variability in these representations independently explain variability in choice.

**Figure 4:**
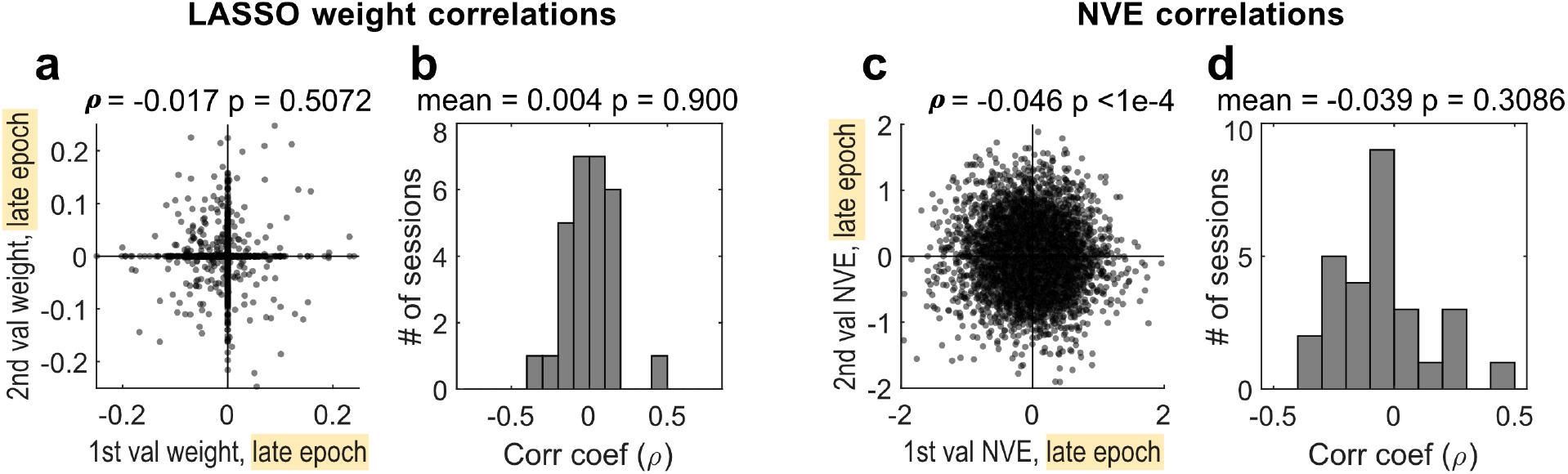
Correlations between LASSO regression weights and between NVEs. **(a)** Correlation between fitted weights from ‘1st value’ and ‘2nd value’ LASSO models (Eqn. 2), using data from the late epoch. Each dot indicates a single cell (n = 1,450 over 32 sessions); note that the many cells have zero weights due to the L1 regularization; results are similar with OLS weights (not shown). **(b)** Histogram of LASSO weight correlations calculated within each session. **(c)** Correlation between NVEs for ‘1st value’ and ‘2nd value’, from the late epoch. On the scatter plot, each dot indicates a single test trial (n = 9,920 over 32 sessions). **(d)** Histogram of NVE correlations calculated within each session.

To determine the fraction of cells that contribute to decoding, the cells with the highest absolute weights in each session were iteratively removed from the model. Classification degraded to near-chance levels after removing about ∼10 cells on average (Fig. 3c, black and blue lines). Decoding accuracy did not depend upon the total number of cells recorded in each session, and was similar for stricter or more liberal criteria for cell sort quality (Extended Data Fig. 7). We conclude that, within the sampling regime of this study, choice outcomes can be decoded from value signals conveyed by a modest fraction of recorded cells (∼20% on average) and that sampling from a significantly larger pool of neurons in the same regions of OFC would not necessarily produce outsized gains in classifier performance.

In contrast to the population-derived decoder, the average predictive accuracy of single cells that contributed to the decoders was negligible. To quantify single-cell predictions, we identified at each time bin the cells with non-zero weights in Eqn. 2, i.e. the subset of cells that contributes to the population-based decoder (For example, for the epoch analyzed in Fig. 2g-j, 599 out of 1450 cells, or a mean of 18.7 per session, had nonzero decoder weights.) For each nonzero-weighted cell we computed single cell choice probabilities (AUCs), with positively-weighted cells assigned first target choices as the positive class, and negatively weighted cells assigned second target choices as the positive class. On average, single-cell choice probabilities of the neurons that contributed to the decoders were at near-chance levels throughout the trial (Fig. 2j and dotted lines in Fig. 3a,b).

Thus, the choice-predictive activity furnished from pooled value signals was not immediately evident at the single cell level. This raises two related questions, addressed below in follow-up analyses. First, what accounts for the superior choice-predictive performance of the population-based decoder? Second, what is the contribution of individual neurons to choice decoding? The superior decoding accuracy of the population model compared to single cells was attributable to the pooling of neuronal activity in single trials. This is shown by a control analysis in which the encoding, decoding, and classification were performed as above, except that the cell weights used for decoding (Eqn. 3) were replaced with either +1 or −1 depending on the sign of the fitted coefficient from Eqn. 2. Under this control, decoding performance was nearly as accurate as the main results (Fig. 3d, “pooled, no weights”). In contrast, a complementary control using the fitted cell weights over unpooled activity produced chance-level decoding (Fig.

To understand the contribution of individual cells to decoder performance, we first examined the single-cell choice probabilities of the neurons that contribute to the population decoder (cells with non-zero weights in Eq. 2.) Although the session-wise mean choice probabilities of these cells was positively correlated with population decoding accuracy (Fig. 5a), on a cell-by-cell basis there was virtually no relationship between the single-cell choice probabilities and the decoder weights (Fig. 5b,c). In other words, single-cell choice probabilities predicted decoder performance in the aggregate, but showed almost no relationship with the weights underlying the decoder’s predictions.

**Figure 5:**
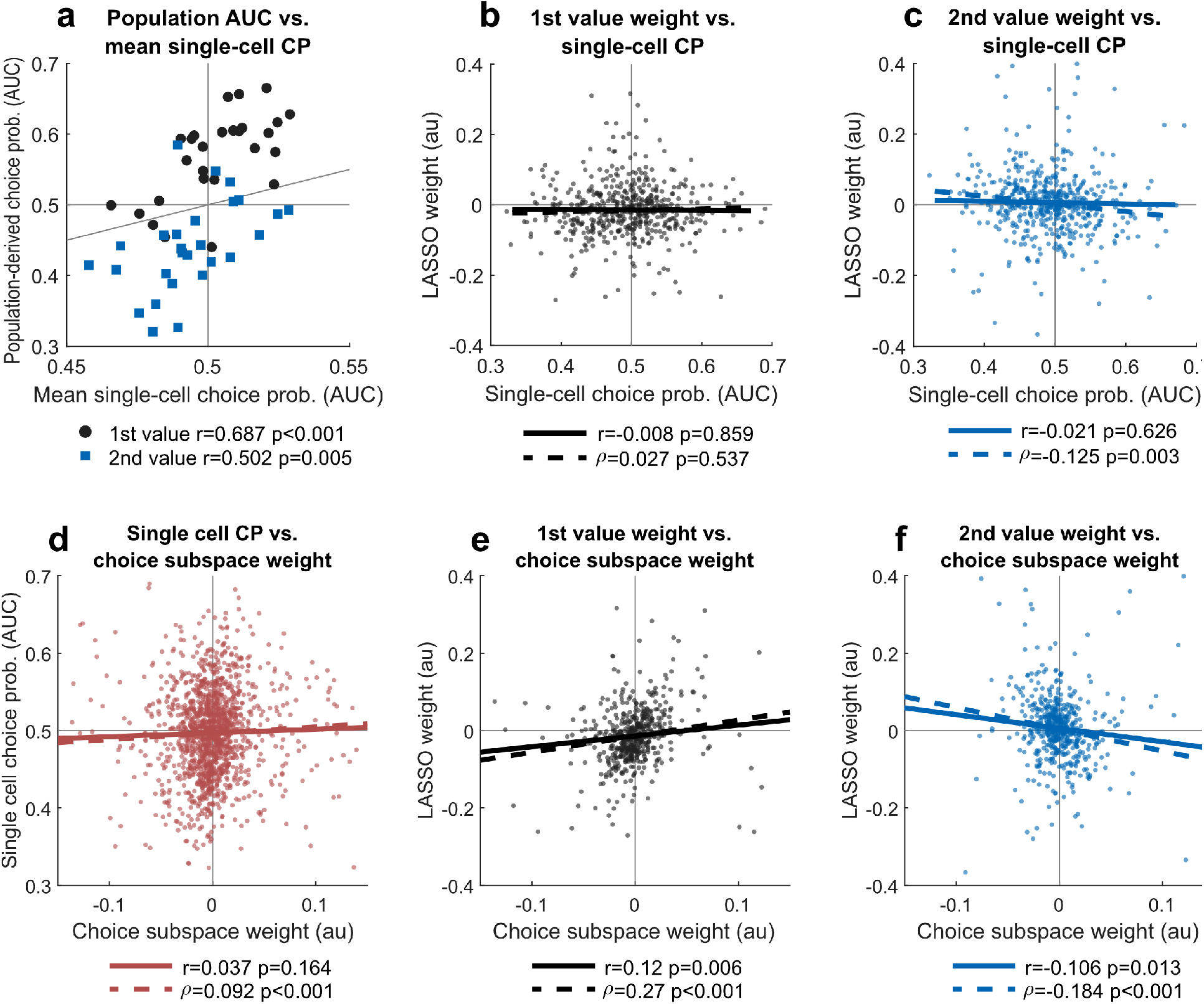
Relationship between value encoding weights, choice subspace weights, and single-cell choice probabilities (CPs). Data are taken from the late epoch in Fig. 3b. Solid lines indicate ordinary least squares fit; dashed lines indicate robust fit from MATLAB function `robustfit` with default arguments. Statistics *r* and *ρ* indicate Pearson’s and Spearman’s coefficients, respectively. **(a)** Session-wise correlation between population-based choice decoding (Fig. 3b.) and the mean single-cell CP of the cells with non-zero decoder weights. Gray diagonal indicates the unity line. **(b-c)** Correlation between single-cell CPs and value-based decoding weights from Eqn. 2. N=531 cells in (b) and n = 551 cells in (c). **(d)** Correlation between single-cell CPs and choice-based decoding weights (n = 1450, see Methods). **(e-f)** Correlations between value-based decoding weights and choice-based decoding weights. N=531 cells in (e) and n = 551 cells in (f). 3d, “weighted, no pooling”). In other words, simply pooling signed neural signals in single trials was sufficient to nearly replicate performance of the full decoder model, whereas preferentially weighting single-cell measures of decoding (choice probability) did not.

One potential explanation for this contradiction is the fact that single cell choice probabilities may be poor estimators of a cell’s behavioral readout (its ultimate contribution to behavior). In particular, theoretical work suggests that given a sufficiently large pool of neurons that contribute to choice, the choice probability of any given cell is a function of the shared noise within the pool rather than that cell’s behavioral contribution^7, 26, 37^. To circumvent these limitations, we took inspiration from recent work in perceptual decision-making; instead of attempting to directly estimate the read out function of single cells, we inferred key properties of the neural read-out mechanism by quantifying the information that can be decoded from the ensemble^8, 38, 39^.

To illustrate this approach, we adopt the neural state-space framework shown in Fig. 6a. The black and blue vectors represent the 1-dimensional subspaces (linear patterns of neural activity) that maximally explain variance in the variables ‘1st value’ and ‘2nd value’, respectively.

**Figure 6:**
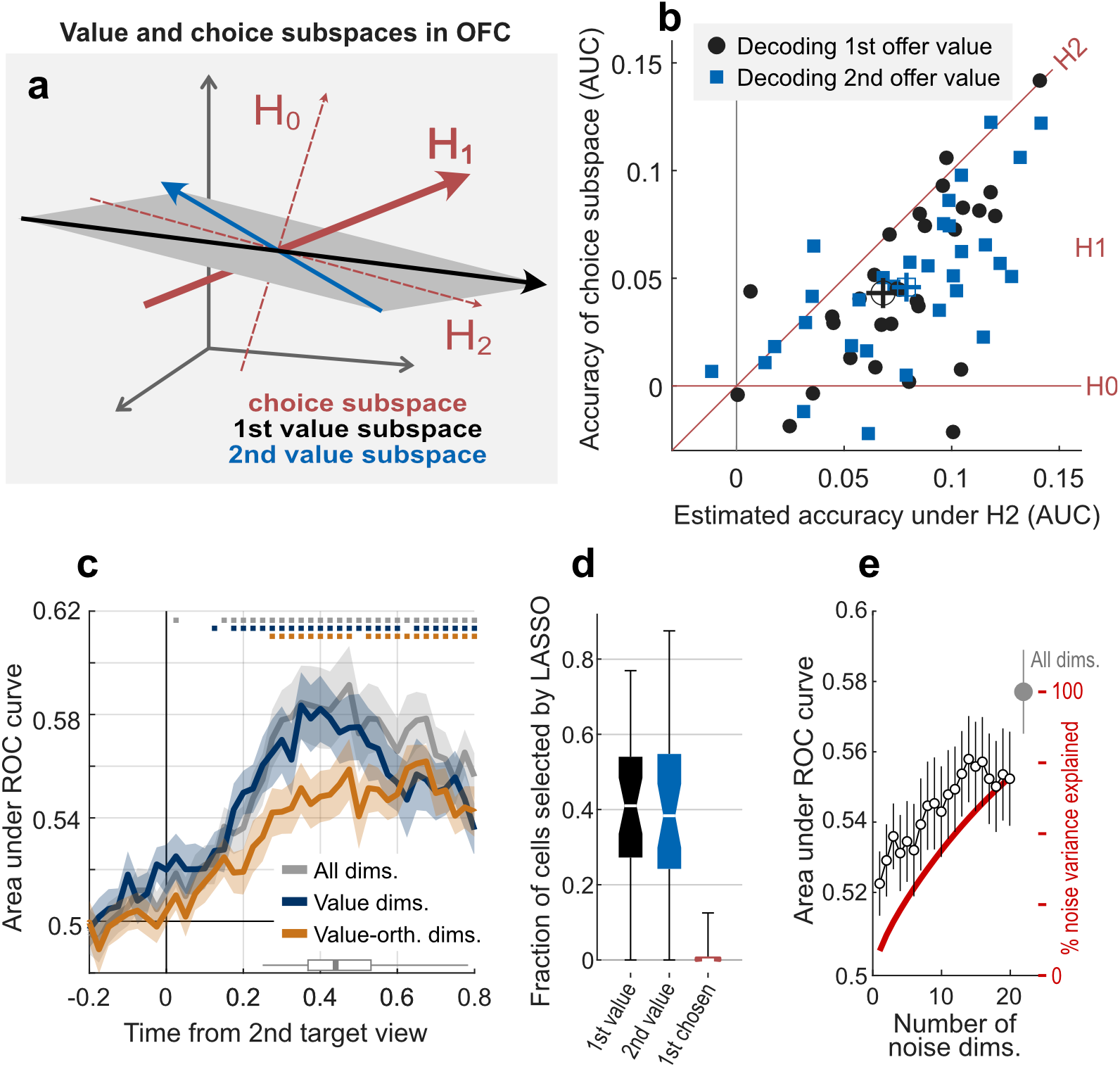
Properties of the OFC choice subspace. Data in this panel are taken from the late epoch in Fig. 3b. **(a)** Illustration of the state space framework used to investigate choice subspace properties. Gray axes indicate a Cartesian space defined by neural firing rates. Black and blue vectors represent 1-D subspaces encoding offer values, and red vectors indicate hypothetical 1-D subspaces encoding choice-predictive activity. **(b)** Decoding offer values within the choice subspace. The y-axis gives the actual accuracy of activity projected into a 1-D choice subspace defined by fits to Eqn. 2, and the x-axis gives the estimated accuracy of an ideal 1-D subspace designed to be contained entirely within the value subspace, as in H2 (see Methods). Dots indicate sessions (n=32), and centroids indicate means and SEM. **(c)** Decoding choice outcomes using the native data (gray), using data projected into the 2-D subspace spanned by the two value vectors (dark blue), or using data projected into the subspace orthogonal to the two value vectors (orange). **(d)** Fraction of cells assigned non-zero weights in LASSO models fit to ‘1st value’, ‘2nd value’, or ‘1st chosen’. Whiskers show the full data range. **(e)** Decoding choice outcomes using data projected into subspaces defined by correlated noise. The x-axis gives the number of noise-defined principal components used for the projection. The y-axis gives the accuracy of choice decoding (left, white dots) and the cumulative percent of noise variance explained by successive components (right, red line).

Together, these vectors define a plane, which we call the value subspace. The three red vectors represent hypothetical subspaces that explain variance in choice – i.e. patterns of activity that may be read out downstream to influence decision outcomes. These three hypothetical subspaces have different relationships with the value subspace: H0 supposes that the choice and value subspaces are orthogonal, and predicts that the choice subspace should contain no information about the offer values. H1 and H2 suppose that the choice and value subspaces are aligned, with H2 representing a choice subspace contained entirely within the value subspace; these predict that the choice subspace should contain nontrivial information about both offer values, with H2 providing the maximum information available from a 1-D subspace.

Recent findings in visual cortex in perceptual choice suggest that the subspaces for choice and sensory inputs are only partially aligned, similar to H1 illustrated in Fig. 6a^39^. To identify the hypothesis most consistent with our OFC data, we reasoned that the degree of alignment between the choice and value subspaces should be evident from the decodability of value information from the choice subspace. To test this, we first removed from the spiking data variance attributable to the offer values by taking the residuals from a factorial ANOVA; we then fit Eqn. 2 to explain the variable ‘1st chosen’, found the resulting fitted weights, took the weighted sum of activity in held-out trials, and used these single-trial choice signals to decode adjacent value levels from those trials (e.g. 2 drops vs. 3 drops) for both the first and second offers. In state space terms, this is equivalent to projecting neural activity into a 1-D choice subspace, and quantifying the information about offer values within this subspace. As shown in Fig. 6b, mean value decoding accuracy was above chance (0 on the y-axis) for both the first and second offer values, leading us to reject H0. To adjudicate between H1 and H2, the decoding under H2 was estimated by decoding values from an “ideal” subspace defined as the vector difference of the weights defining the ‘1st value’ and ‘2nd value’ subspaces (see Methods). Value decoding accuracy from the actual choice subspace was below that of the ideal subspace, indicated by the scatter of points below the unity line in Fig. 6b. These results favor H1.

To confirm H1 we used a complementary method in which we decoded choice outcomes from neural activity projected only into the value subspace (the plane in Fig. 6a), or only into dimensions orthogonal to the value subspace. Data projected into the value-only subspace furnished decoding accuracies essentially equivalent to those obtained when using all dimensions (dark blue vs. gray in Fig. 6c), whereas the value-orthogonal subspace showed lower, but still above-chance choice decoding accuracy (orange in Fig. 6c). In other words, the OFC contains both value- and non-value-related patterns of neural activity that predict choice. In contrast, decodable representations of offer values appear to be confined entirely to a 2-D value subspace (Extended Data Fig. 4). Control analyses show that the results are no different when chosen value signals are first removed (Extended Data Fig. 8), and that predictive activity in an order-based reference frame is more accurate than in a spatial reference frame (Extended Data Fig. 3d). The results in Fig. 6c therefore suggest a more nuanced version of H1, in which the OFC contains multiple patterns of choice-predictive activity, and in which value signals are contained within a low-D subspace that projects onto a highly choice-predictive subset of choice-related patterns.

Returning to the question of individual cell contributions, the nontrivial alignment between the value and choice subspaces suggests that at the single cell level there should be an association between representations of value and choice. Consistent with this, we found that value decoder weights were significantly correlated with the choice subspace weights (Fig. 5e,f). In addition, the choice subspace weights were only weakly correlated with single-cell choice probabilities (Fig. 5d), consistent with the idea that single-cell choice probabilities are poor estimators of a given cell’s contribution to choice^7, 26, 37^. This suggests that the choice subspace weights (derived by joint estimation in Eqn. 2) are a more accurate reflection of a cell’s contribution to a decision compared to the single-cell choice probabilities. One potential limitation, however, is that behavioral read-out weights may be difficult to accurately estimate from small samples of cells, as illustrated in computational work by Ruff et al.^8^.

Taking the choice subspace weights as an index of contribution to behavior, we also found that single cells with strong choice-modulated activity were rare compared to those that strongly encoded value. Specifically, using LASSO regularization in Eqn. 2 to estimate the choice subspace produced model fits in which only ∼1% of cells had non-zero weights; in contrast, ∼40% of cells were assigned non-zero weights for LASSO models fit to ‘1st value’ or ‘2nd value’. (Fig. 6d). Therefore, unlike offer values, the neural coding of choice in OFC appears to be defined by weak firing fluctuations distributed over many neurons, rather than by strong changes in firing rate in a subset.

Finally, the downstream read out of neural activity can be limited by the magnitude and structure of correlated noise in the population, especially when the activity patterns that characterize shared noise overlap with activity patterns related to choice or stimulus encoding ^27, 40, 41^. The mean noise correlation for all cell pairs was 0.019 (STD 0.083, n=39,049 pairs), virtually identical to the mean reported by Conen and Padoa-Schioppa (2015); among nonzero-weighted cells, the noise correlations were smaller (mean 0.013, STD 0.081, n=2,579 pairs, see Extended Data Fig. 9). In visual area V4, the choice subspace is strongly aligned with the first principal component of shared variability in the population^39^, indicating suboptimal read-out of motion information for choice. In contrast, in our OFC data choice decoding accuracy within the first noise principal component was low, and increased steadily as a function of the number of shared noise dimensions used (Fig. 6e). The overall low noise correlations and lack of strong alignment between the noise and choice subspaces suggests that, unlike in V4^39^, the behavioral read out from OFC is not substantially limited by shared variability.

## DISCUSSION

In summary, we identify for the first time a robust trial-to-trial relationship between variability in OFC value representations and variability in value-based choices in primates performing at behavioral threshold. This newly identified neural-behavioral link represents the culmination of important prior work that links OFC value representations to decision reaction times^12^ and to choice behavior at the session level^11^, as well as detailed analyses of single-cell choice probability effects in different OFC sub-populations^23, 25^. In addition, we identify for the first time key properties of the OFC choice subspace, defined as the multi-neuron activity patterns that explain variance in choice outcome independent of value. These findings are significant because they fill a critical missing piece of a long-standing puzzle in neuroeconomics – i.e. they show how value representations realized at the level of neural spiking activity contribute to the outcome of value-based choices at the resolution of single trials. In addition, the identification of the OFC choice subspace and its properties gives new insights into the general mechanisms by which OFC activity is read out downstream.

The novel findings were made possible by two critical design elements. The first was a large fraction of equal-value choice conditions, which elicited variable choices and provided maximum leverage for detecting a relationship between choice outcome and neural activity. The second was ensemble recordings and analyses, which permitted the pooling of neural activity in single trials, and therefore the single-trial estimation of population value representations. While these analyses also involved the selective weighting of cell activity, the bulk of the choice-predictive effect can be explained by single-trial pooling without selective weights, (see Fig. 3d).

Taken together, our findings suggest a parsimonious mechanism for how value representations in OFC are linked to behavior: projection of value information onto an output-potent subspace. The term output-potent subspace refers to the idea that within a given neural population, only certain activity patterns are effective for driving the activity of downstream regions^38, 42, 43^. Likewise, the representation of offer values can be conceived of as input subspace, patterns of neural activity that are driven by feedforward sensory inputs as the stimuli are sampled during the trial, and that represent behaviorally relevant stimulus information (reward association). Our findings show partial alignment between the input and output subspaces in OFC. In other words, they suggest a mechanism in which neural activity patterns that encode values overlap with activity patterns that are read out downstream and that consequently influence choice. Large fluctuations in value signals (e.g. for high vs. low value offers) result in large displacements within the choice subspace and therefore large differences in the probability of choosing the first offer; and small fluctuations (e.g. due to neuronal noise across trials of the same type) correspond to smaller displacements in projected activity, and to smaller variations in decision probabilities that are most apparent for near- or equal-value offers. In this way, representations of value are mapped onto choices.

An important limitation of the current study is that the characteristics of the putative read out of OFC activity are inferred based only on correlational evidence – the trial-wise relationship population-level neural activity and choice. To fully test the mechanism proposed above will require manipulation of OFC activity within the value and choice-encoding subspaces, possibly by selective manipulation of individual neurons whose activity defines these subspaces^44^.

Another limitation is the simplifying assumption that the offer values are represented by linear, 1-D subspaces. While the results in Extended Data Fig. 4 are consistent with this assumption, recent findings in vmPFC suggest curvilinear value encoding requiring multiple dimensions to fully explain^45^. A third limitation is the fact that the subspace alignment described in Fig. 6 pertains to a single time point following the second offer; however value subspaces may be dynamic from one moment in time to the next^22, 29, 46^. To identify whether choice subspaces are also dynamic will require prolonging the decision time course to allow for comparison of the choice subspace across epochs. Finally, the generalizability of the proposed mechanisms is currently unknown. For example, supplementary analyses identified spatial-based value and choice signals that appear to have only a secondary role in this paradigm (Extended Data Fig. 3); however, in other cognitive tasks, OFC spatial signals may play a more prominent role (e.g. Yoo et al. 2018^47^).

While prior studies give different perspectives on how values are represented in OFC ^12, 22, 25, 29, 48^, the present findings offer a framework for understanding the critical question of how these representations are ultimately read out – a framework that is consistent with observations in other cortical areas and other behavioral domains, and that may therefore reflect general principles of cortical information flow^39, 42, 43^. In this way, the present findings provide a path forward for adjudicating between and unifying the diverse perspectives on the function of primate OFC.

In the illustration in Fig. 6a, both the choice subspace and value subspaces are shown as 1- dimensional vectors; likewise, the decoding approach used assigns only one weight to each cell, and therefore considers only the information available along a single dimension in each model. However, our results may be inconsistent with this simple geometric interpretation of the choice subspace. If the choice subspace were strictly 1-dimensional (i.e. the vector H1), and if decoding accuracy reflects the magnitude of the projection of this vector onto the value subspace, then the decoding accuracy available from all dimensions (gray) should be the vector sum of the accuracies in the value and value-orthogonal subspaces (dark blue and orange). However, this is not the case: Fig. 6c shows that the accuracy within the value dimensions is essentially equal to that available from all dimensions – and yet the value-orthogonal dims also show substantial accuracy as well. In other words, it suggests that there are many potential patterns of OFC activity that can modulate decision behavior, that these patterns are not fully captured by a 1-D vector, and that representations of the stimulus values (which in the present study appear to be low-dimensional, Extended Data Fig. 4) access a limited, yet still highly-predictive subset of these patterns.

Decisions in the natural world often entail weighting multiple attributes, such as magnitude, probability, availability, etc., and in this regard a multidimensional choice subspace in OFC would permit many decision features to simultaneously influence behavioral outputs without destructive interference. Likewise, it has been proposed that portions of OFC are specialized for constructing value with reference to specific classes of objects or object features^49, 50^. Given the observation that OFC value signals can predict choice outcomes, a logical follow-up question is whether these predictions are best explained as a function of “pure” value signals that reflect only the expected utility of a given offer; or whether they are best explained by a combination of feature-specific value representations. To answer this question would require new experiments, e.g. that manipulate the mapping between offer attributes and the physical features of choice stimuli. At present, our results are compatible with either mechanism.

Implicit in the mechanism proposed above is the idea that the encoding of choice outcomes in OFC does not reflect an explicit feedforward input or the outcome of a locally mediated computation; rather, the encoding of choice outcomes emerges as a consequence of the OFC’s outputs to downstream regions involved in the decision process. It also implies that the comparison between offers does not necessarily take place in the OFC, but instead takes place downstream. In contrast, the offer values are explicitly represented, in the sense that feedforward visual inputs drive the activity of OFC neurons in a manner that depends on the reward association of those stimuli. Consistent with this, regularized regression identifies a large fraction of OFC neurons that explain variance in offer values, whereas very few individual neurons are identified as explaining variance in choice independent of value (Fig. 6d).

Consistent with prior work, the average noise correlation between pairs of OFC neurons was ∼0.019, which is roughly 6 times smaller than the correlations observed in area MT, and is smaller than in many other cortical areas as well (see Figure 3a in Conen & Padoa-Schioppa 2015). Correlated noise plays a critical and complex role in shaping the information that can be decoded from single neurons and neural populations^41^. Most relevant to the present study is the fact that net positive noise correlations permit the information encoded by a large population of neurons to be inferred using only small sample of cells^8^. Moreover, given a sufficiently large population of neurons carrying decision-relevant signals, the choice probability measured in any given cell is almost entirely attributable to its covariance with other cells in the population, rather than its fractional influence on choice^7, 26, 37^. Therefore, the lower noise correlations observed in OFC are entirely consistent with the average magnitude of choice probabilities in single cells (as explored in depth by Conen & Padoa-Schioppa^25^), and by extension explain the relatively modest population-derived decoding accuracy observed in OFC, as compared to regions with higher noise correlations, such as MT^3^.

The net effect of shared variability on neural information transmission is also a complex issue (see Kohn et al.^41^ for an in-depth discussion). One relevant factor is whether population-level patterns of correlated noise are aligned with the activity patterns that carry choice-relevant information^40^. For example, within populations of simultaneously recorded visual area V4 neurons, Ni et al.^39^ showed that the first principal component defined over the noise correlations captured essentially all of the decision-relevant neural variability in the population, indicating strong alignment between noise and choice-related activity patterns. This has been interpreted as reflecting a “general decoding” mechanism, in which neurons share variability along all of the neural dimensions that encode potentially relevant features (e.g. color, orientation, binocular disparity), rather than just the dimensions relevant to a single task. In contrast, in OFC we observed that noise-defined subspaces did not capture an outsized portion of decision-relevant neural activity (Fig. 6e). While more work is necessary to better understand this finding, one interpretation is that the lack of strong alignment between noise- and choice-related activity may be due to the much greater flexibility of OFC neural coding compared to visual areas^29, 48, 51^. According to Ni et al., the alignment between noise and choice-related activity in V4 emerges as a consequence of V4 having relatively static read-out weights for behavior. In contrast, if the behaviorally-potent subspace of OFC is dynamic, then the shared noise would not be expected to strongly align with choice-related activity in any given task or context.

## METHODS

### SUBJECTS AND APPARATUS

All procedures were performed in accordance with the NIH Guide for the Care and Use of Laboratory Animals, and were approved by the Animal Care and Use Committees of Stanford University, and Rutgers University – Newark. The subjects were two adult male rhesus macaques designated K and C, weighing ∼14kg each at the time of the study. Data from Monkey K were acquired at Stanford University, and from Monkey C were acquired at Rutgers University – Newark. They were implanted with MR-compatible head holders and recording chamber (Crist Instruments, Hagerstown, MD); craniotomies were also performed to allow access to the OFC. Monkey K had bilateral chambers and craniotomies, permitting bilateral recordings. Monkey C had a single chamber and craniotomy overlying the left OFC. All procedures were performed under full surgical anesthesia using aseptic techniques and instruments, with analgesics and antibiotics given pre-, intra- and/or post-operatively as appropriate. Data were collected while the monkeys were head-restrained and seated ∼57 cm from a fronto-parallel CRT monitor displaying the task stimuli (120Hz refresh rate, 1024x768 resolution). Three response levers (ENV-612M, Med Associates, Inc., St. Albans, VT) were placed in front of the subjects, within their reach. The “center” lever was located approximately 21cm below the display center, and the other two levers were located approximately 8.5cm to the left and right of the center lever. Stimulus presentation, reward delivery, and monitoring of behavioral responses was controlled through MATLAB (Mathworks, Inc., Natick, MA), using the ROME toolbox (R. Kiani, 2012). Horizontal and vertical eye position was recorded noninvasively at 250Hz (Eyelink, SR Research, Mississauga, Ontario, Canada). Fluid rewards were delivered via a gravity-fed reservoir and solenoid valve. Neural activity, eye position, and task event data were acquired and stored using a Plexon Omniplex system (Plexon, Inc., Dallas, TX). Analysis was performed using custom routines in MATLAB 2019b and the R computing environment (version 4.0.2) with the *glmnet* package (version 4.0-2).

### DECISION TASK

#### Task structure

Fig. 1a shows an abbreviated task sequence; Extended Data Fig. 1a shows the full sequence as described here: Monkeys initiated a trial by fixating a central point on the display, and manually depressing the center lever. After holding fixation and the center lever for a variable duration of 1-1.5s, the fixation point disappeared, and two target arrays appeared, centered 7.5 degrees of visual angle to the left and right of the display center (Fig. 1a). As soon as the arrays appeared, the monkeys were allowed to shift their gaze as they desired to view the targets, in any order and for any amount of time, until they initiated a choice. Eye position was monitored throughout, but had no programmatic effect on the trial outcome (with the exception of enforced central fixation necessary to initiate a trial). The monkeys were encouraged to view each target at least once, through the use of visual crowding and gaze contingent programming, described in detail below. The monkeys were free to choose at any time, by first lifting the center lever (extinguishing the targets), and then pressing either the left or right lever to choose the left or right target, respectively. After the left/right press, a juice reward of 1-5 drops was delivered, depending on the value of the chosen target (Fig. 1b). To discourage the monkey from deliberating or changing their minds after the center lever lift, the interval between center lift and left/right press was limited to 400ms (Monkey K) or 500ms (Monkey C); failure to respond within this deadline resulted in an aborted trial.

#### Choice targets

The choice targets were colored glyphs, each a unique combination of one of four colors and one of four shapes. A full example set of 12 targets is shown in Fig. 1b. New targets sets were generated every 1-5 sessions, to minimize over-learning of the stimuli. The available shape primitives and the set generation procedure is given in Extended Data Fig. 1b.

Offer values ranged from 1-5 drops of juice. Importantly, every value stratum contained at least two unique targets, permitting us to offer choices between two distinct but equal-value targets. Note that the terms ‘offer value’ and ‘target value’ refer to a target’s reward association (1-5 juice drops), and the term ‘target identity’ refers to a target’s unique color/shape combination, of which there were a total of 12 in every set (Fig. 1b). Targets and crowders (see below) were all 0.75x0.75 degrees of visual angle. The patch colors used for targets and crowders were selected to be isoluminant. The nominal patch luminance for all colors used in Monkey C’s sessions was ∼2.7 cd/m^2 with negligible background luminance; and for Monkey K’s session was ∼22 cd/m^2 with background luminance of ∼4cd/m^2, as measured by a Tektronix luminance J17 photometer with a J1820 head. Note that during data collection, the luminance of the target stimuli was reduced by up to ∼50% of these nominal values (with no change to the crowders), in order to better obscure the targets within the crowder array (Fig. 1c). This luminance reduction did not prevent the monkeys from identifying the targets, because choice performance was nearly perfect for the easiest trials (Fig. 1d, value differences of 3 and 4).

#### Target arrays, visual crowding, and gaze-contingent mask

Two methods were used to encourage the monkeys to directly fixate both targets prior to making a choice. First, each target was surrounded by six ‘crowders’, which were randomly generated, multi-colored ‘+’ or ‘×’ shapes that had no direct association with reward. (Fig. 1c). Crowders reduce the effectiveness of peripheral vision^30, 31^, making it difficult to identify a target without using high-acuity foveal vision. Second, we used gaze-contingent programming to initially obscure the targets until the first eye movement was made: At the moment the target arrays first appeared on the display, the two targets were masked with a randomly generated crowder (Extended Data Fig. 1a). Therefore, no information about target identity or value was initially available. At the moment the monkey’s eye position breached the central fixation window (diameter 3.5-4 degrees), both masks were removed, and the choice targets were shown in their place, one each at the center of the two target arrays. Because the display frame rate interval (∼8ms) was shorter than the time needed to complete a saccade (∼20-40ms), the masks were usually replaced with the targets while the eyes were still moving. Thus, the task design ensured that the monkeys could gain no information about the offer values until they initiated a saccade towards one of the two targets. After the targets were shown, they remained in place until the monkey initiated a choice by lifting the center lever, at which point both targets and all crowders were extinguished.

### NEURAL RECORDINGS

Linear recording arrays (Plexon V-Probes) were introduced into the brain through a sharpened guide tube whose tip was inserted 1-3mm below the dura. Probes had 16, 24, or 32 channels, spaced either 50 or 100µm apart; up to 2 probes were used per hemisphere (32-80 channels, mean 50 for monkey C, 75 for Monkey K). OFC was identified on the basis of gray/white matter transitions, and by consulting a high-resolution MRI acquired from each animal. We targeted the fundus and lateral bank of the medial orbital sulcus and the laterally adjacent gyrus (Extended Data Fig. 2), over anterior-posterior coordinates ranging from +32 to +38mm anterior to the intra-aural landmark^52^.

Neural unit signals were sorted by automated spike detection and sorting routines (Kilosort2, default parameters, except for ops.th = [4 12] and ops.lam = 15) followed by manual curation (Phy2 GUI). Sorted units were considered eligible for analysis if they had less than 2% of inter-spike intervals (ISIs) below 2ms, and also had less than 25% of spikes estimated to have come from ‘rogue’ units, according to the methods of Hill et al^53^. Based upon follow-up manual inspection (Plexon Offline Sorter), these criteria selected all well-isolated single units, as well as less-well isolated units lightly contaminated with rogue spikes. We collectively refer to units that meet these thresholds as ‘cells’, recognizing that they represent a mix of units of varying isolation quality. The key findings in this study were confirmed when using a more stringent 1% ISIs threshold, and 10% rogue spike threshold (Extended Data Fig. 7).

### ANALYSIS

#### General statistical procedures

Data from the two animals were combined for all analyses, except in Extended Data Fig. 5a-h. Hypothesis testing for group means was performed using two-sided t-tests, unless the underlying data were skewed, in which case Wilcoxon sign-rank tests were used. Statistical association was measured using either Pearson’s *r*, Spearman’s *ρ*, or both. Boxplots give the median and inter-quartile range, and each whisker extends to 1.5 times the interquartile range. Time series data show the mean and SEM (shaded region). Bar graphs and error bars show mean and SEM, unless otherwise indicated. Significance in time-series graphs is indicated at p<0.05 corrected for multiple comparisons within each time series, using Holm’s Bonferroni method. Unless otherwise stated, all other p-values were uncorrected.

#### Behavior and key task variables

Choice performance was quantified by a logistic regression in which the fraction of left choices was explained by the left-minus-right offer value (in drops of juice, Fig. 1d). The number of fixations per trial and the decision RTs were averaged by task difficulty, defined as the best-minus-worst offer value (Fig. 1e,f). For neural data analyses, only the trials in which monkeys viewed both targets were used (99.3% of trials for Monkey C, and 93.2% of trials for Monkey K); among these, the vast majority of trials had either two or three total fixations. Consistent with similar tasks performed by humans^54^, the monkeys showed a strong tendency to devote their final fixation onto the item they chose (for details see Lupkin & McGinty, 2022^55^). This means that trials with exactly two fixations (e.g. Left then Right) were very often trials in which the second-viewed item was chosen, and trials with exactly three fixations (e.g. Left then Right then Left again) very often ended in choices in favor of the first-viewed item. Because of the association between total fixations and choice, we did not separately analyze two- and three-fixation trials, but instead combined together all trials with two or more total fixations.

Neural data analysis was based on three task variables defined in a viewing order-based reference frame: ‘1st value’ and ‘2nd value’ were defined as the number of juice drops associated with the first and second target (respectively) viewed in every trial; and ‘1st chosen’ was defined as *true* when the monkey chose the first target he viewed in the trial, and *false* otherwise. For single-cell analyses, we also defined ‘chosen value’ as the value of the offer chosen on each trial.

#### Single cell linear model and coefficient of partial determination (CPD)

Data were time-locked to the first and second offer viewing times, with 200ms bins and 25ms increments. To quantify the encoding of the task variables, we first fit the following ordinary least squares linear model for each cell at every time bin:

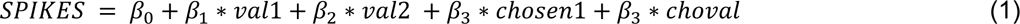

Where *SPIKES* is the spike count of a single cell in a single time bin, *val1, val2, chosen1,* and *choval* are the variables ‘1st value’, ‘2nd value’, ‘1st chosen’, and ‘chosen value’ as defined above. Encoding was quantified by computing for each variable the coefficient of partial determination (CPD)^22^, which captures the unique variance explained by each variable after accounting for the variance explained by all the others. Within each time bin, the mean and SEM of the CPD was calculated across cells; the results are shown in Fig. 2d,e.

#### Noise correlation

Pairwise Pearson’s noise correlations were measured from spike counts over a 200ms window centered 200ms after fixation onto the first target. Before calculation of the noise correlations, spike counts were first z-scored according to the trial condition, defined as the identity of the first target and whether the first fixation location was left or right.

#### Population-based decoding

At a high level, this approach entails the classification of choices offer values, using neural signals derived from the pooled activity of many simultaneously recorded neurons. The primary analysis entails decoding choice outcomes from population-derived value signals (results in Figs. 2g-i, and Extended Data Figs. 5 and 7). Additional analyses entail the reverse: decoding offer values from population representations of choice outcome (results in Fig. 6b). Finally, results in Extended Data Fig. 4 show decoding offer values from population-derived value signals, and results in Fig. 6c,e and Extended Data Figs. 3d and 8b show decoding of choice outcomes from population-derived choice signals. Classification was performed with receiver operating characteristic analysis (ROC) to facilitate comparison to choice probability metrics in prior studies ^1, 23–25^. Population-level value signals were obtained using an L1-regularized regression model (LASSO)^35^, although similar results were obtained using Ridge or ordinary least squares (OLS, not shown).

Decoding proceeded in three steps: fitting a model to explain offer value as a function of neural activity (encoding); using the model to obtain a neurally-derived estimate of offer value in held-out trials (decoding); and classification of choice outcomes using the decoded value estimates (classification). Note that the encoding and decoding steps can be interpreted as a form of dimensionality reduction and projection: the fitted parameters of the encoding model define an axis (a 1-D subspace) within the neural state space that encodes value, and the decoded estimates quantify the projection of neural activity onto that axis.

This analysis used spike counts observed in 200ms bins with 25ms increments, time locked to the viewing of the first and second offer in each trial. Results were similar when using 100ms bins (not shown). These steps were performed independently within each bin, meaning that fitted parameters could vary across bins – in particular, that from one bin to the next, different cells could be assigned non-zero weights in the LASSO model described in Eqn. 2 below. For some follow-up analyses we defined an ‘early’ epoch, 0.2 to 0.4s after first target viewing; and a ‘late’ epoch, 0.2 to 0.4s after second target viewing (Fig. 3a,b).

First, in the encoding step, the data were stratified into a training and test set, maintaining whenever possible equal numbers of trial types, defined by the values of the first- and second-viewed offer in each trial. Then we fit the following L1-regularized linear model (LASSO)^35^ to the training trials, using the `glmnet` package in R:

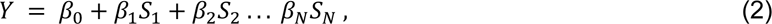

where 𝑌 was either ‘1st value’ or ‘2nd value’, 𝑆_𝑖_ was the spike count observed from cell *i* in a given bin, and *N* was the number of simultaneously recorded cells in a session. The resulting fitted parameters, {𝛽_1_, 𝛽_2_, . . . 𝛽_𝑁_}, can be interpreted as a set of weights that indicate the degree to which the firing of each cell uniquely and monotonically contributes to explaining variance in the dependent variable 𝑌. The model was fit by minimizing the negative log likelihood plus a penalty 𝜆|𝛽_2_, . . . 𝛽_𝑁_}|_1_, where ||_1_ indicates the L1 norm. The scaling parameter 𝜆 (lower-bounded by zero) was selected to minimize cross-validation error (10-folds) within the training trials^35^.

This form of regularization results in a sparse set of weights^35^, such that the weights for cells that explain negligible variance are set to zero. In this way, the model unambiguously selects a subset of cells (those with non-zero weights) whose activity contributes to the encoding of value. Having a clearly defined subset of cells is important, because it allows the population-based choice decoding effects to be directly compared to single-cell choice probabilities over the exact same cells (see below).

Before fitting Eqn. 2, the effects of the non-relevant value variable were removed from the spiking data (training trials only). When 𝑌 was the variable ‘1st value’, the spiking data were normalized by mean-centering the spike counts with respect to the second target identity. Likewise, when 𝑌 was the variable ‘2nd value’, the spiking data were mean-centered according to the first target identity. Next, in the decoding step, we computed the following weighted sum in a set of held-out test trials:

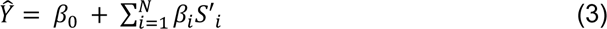

where 𝛽_0_ and 𝛽_𝑖_ were the intercept and fitted parameters from Eqn. 2, 𝑆′_𝑖_ was the spike count from the *i-*th cell, and *N* was the total number of cells. This yields for every held-out trial the weighted sum of spike counts across all simultaneously observed cells. In their raw form, these weighted sums estimate the objective variable 𝑌 in Eqn. 2; for example, when the objective variable was ‘1st value’, 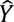 was on average a monotonically increasing function of the first-viewed offer value (Fig. 2g.)

To assess the accuracy of the decoder model for estimating offer values, 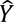 was used to classify trials with adjacent-value offers (e.g. 3 drops vs. 4 drops) using receiver-operating characteristic (ROC) analysis over held out test trials; see Extended Data Fig. 4 for details.

To assess the accuracy of the decoder for predicting choice outcomes (independent of offer values), we consider the trial-to-trial variability in 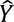, i.e. the degree to which the decoded estimates exceed or fall short of their expected value from one trial to the next. To isolate this variability, 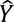 was normalized by taking the residuals from a two-way factorial ANOVA, in which 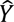 was explained by the both identity of the first-viewed target and the identity of the second-viewed target (with no interaction term). This normalized 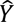 is referred to as the Normalized Value Estimate (NVE, see Fig. 2h). Note that normalization was performed with respect to the 12 unique target identities, rather than to the 5 levels of reward value (Fig. 1b.); this was necessary to control for the possibility that the monkeys may idiosyncratically assign different subjective values to targets that have equal reward associations.

The NVEs therefore reflect trial-to-trial variability in the population-level representations of the offer values. The NVEs were used to classify held-out trials according to whether the first target was chosen, using standard receiver-operating characteristic (ROC) analyses. While encoding and decoding were performed over all of the training and test trials (respectively), classification was only performed over test trials for which the two targets differed by 0 or 1 drop of juice, because these were the only trials with sufficient choice variability (mean 145 trials per session for Monkey K; 175 trials per session for Monkey C). This produced for every session one area under the ROC curve (AUC) for the ‘1st value’ model and one for the ‘2nd value model’. The positive class was always defined as choices in favor of the first-viewed offer. Therefore, the AUC was above 0.5 when higher NVEs corresponded to a greater tendency to select the first target, and was below 0.5 when higher NVEs were associated with a tendency to select the second target.

For the analyses shown in Fig. 6, Eqn. 2 was fit with Y set to the variable ‘1st chosen’, using spiking data that was residualized with respect to the target identities using a two-way factorial ANOVA in which spiking was explained by both the identity of the first-viewed target and the identity of the second-viewed target, with no interaction term. Ordinary least squares was used rather than LASSO, because LASSO fits typically returned non-zero weights for only ∼1% of cells (Fig. 6d). Decoding and classification of offer values (Fig. 6b) or choice outcomes (Fig. 6c,e) in held out trials proceeds as above.

#### Subspace projections

To project the spiking data into different subspaces, we used the method of Semedo et al.^42^. To project the data into the value encoding subspace, we first obtain the fitted weights from Eqn. 2 when 𝑌 was ‘1st value’ and ‘2nd value’, and concatenated these two vectors into a n-by-2 matrix ***B***, where n is the number of cells in a session. The two columns of ***B*** therefore define the value coding subspace – the patterns of neural activity that together explain maximal variance in ‘1st value’ and ‘2nd value’. To find an orthonormal basis for ***B***, we compute ***M*** = ***B***^T^*cov(***X***), where ***X*** is a number-of-trials-by-n matrix of spike counts, and cov(***X***) is the covariance matrix of ***X***. We then find the singular value decomposition ***M*** = ***UDV***^T^. Let ***Q*** be the first two columns matrix ***V***, corresponding to the non-zero singular values of the decomposition, which we use as the projection matrix. Specifically, we find 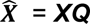, which projects the spiking data X into the subspace spanned by the columns of B – i.e. the plane defined by the encoding vectors for ‘1st value’ and ‘2nd value’. We then perform the encoding, decoding, and classification on 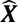. To project data into the subspace orthogonal to ***B***, we find the projection matrix ***P*** by taking all except the first two columns of V, corresponding to the zero singular values of the decomposition, and perform the decoding analysis using data projected into the ***P*** subspace 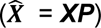.

To obtain subspaces defined by correlated noise, spiking data were sampled from a 200ms window centered 200ms after fixation onto the first target, z-scored according to the identity of the first target and whether the first fixation location was left or right, and then arranged into a number-of-trials-by-n matrix. The PCA loadings from this matrix were taken as columns of the projection matrix, up to the desired number of dimensions. The spiking data ***X*** were then projected into the PCA-defined subspace; the projected data were then used for encoding, decoding, and choice classification as above.

#### Choice probability in single cells

This analysis used the same binned spiking data as the population-based choice probability analysis described above, except that it was performed individually for each cell. Before analysis, spike counts that were first normalized with respect to the target identities, using a two-way factorial ANOVA in which spiking was explained by both the identity of the first-viewed target and the identity of the second-viewed target, with no interaction term. This is the same normalization used to transform decoded value signals (obtained from Eqn. 3) into NVEs. ROC calculations were based upon those trials in the test set in which the offer values differed by 0 or 1.

To directly compare the single cell with population results, average choice probabilities were calculated for two non-exclusive groups of cells: cells with non-zero weights in Eqn. 2 when 𝑌 was ‘1st value’; and cells with non-zero weights when 𝑌 was ‘2nd value’. In other words, mean choice probabilities were calculated for the cells that contributed to the NVE calculations for each respective variable. The positive class of each cell was determined by the sign of the weight from Eqn. 2: when the weight was above zero, the positive class was defined as choices in favor of the first-viewed target; and when the weight was below zero, the positive class was defined as second target choices. This ensures that the direction of the single-cell ROC results (i.e. whether the AUC is above or below 0.5) have the same interpretation as the population-based ROC results.

#### Neuron dropping

The population decoding analysis above was repeated over a successively smaller pool of cells, using data from the late epoch in Fig. 3b. Within a session the *N* recorded cells were ranked according to the absolute value of the weights obtained in Eqn. 2. Then, the cell with the highest absolute weight was removed from the pool, and decoding and classification steps were performed, yielding an AUC based upon the *N-1* remaining cells. This procedure was repeated until no cells remained. Results are plotted for the first 20 removed cells.

#### Logistic regression with behavioral and neural regressors

The fraction of first target choices in the test trials was explained by the following logistic regression:

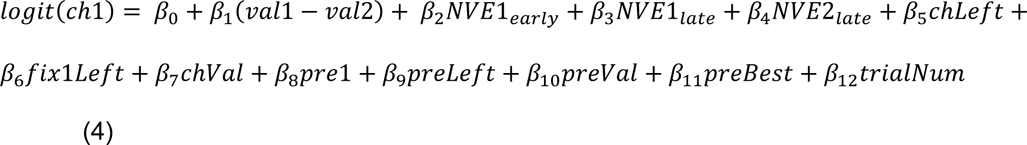

where logit() is the logit function, *ch1* is a Boolean variable indicating a choice in favor of the first-viewed target, *val1* and *val2* are variables ‘1st value’ and ‘2nd value’ defined above, *NVE1* and *NVE2* refer to the NVEs for ‘1st value’ and ‘2nd value’ defined above, and the subscripts indicate data measured at the ‘early’ and ‘late’ epochs indicated in Fig. 3a,b. The model also includes nuisance variables that capture behavior on the current and previous trial: ‘*chLeft*’ indicates whether the left item was chosen on the current trial; ‘*fix1Left’* indicates whether the first fixation was to the left item on the current trial; ‘*pre1*’, ‘*preLeft*’, and ‘*preBest*’ are boolean variables indicating whether the previous trial’s choice was in favor of the first item, in favor of the left item, or was in favor of the highest value item, respectively; ‘*preVal*’ gives the value of the previously chosen offer; ‘*trialNum’* gives the current trial number. The model was fit for each session, and then the regression estimates were averaged across sessions. Non-Boolean variables were z-scored before fitting.

## Acknowledgements

W.T. Newsome for funding, material support, and thoughtful discussions; A. Rangel for thoughtful discussions; J. Brown, E. Carson, S. Fong, A. McCormick, M. Ortiz, J. Powell, J. Sanders, D. Siegel for technical assistance; and J. Corbo, R. Ebitz, D. Headley, G. Karpov, D. Kimmel, M. Rosenberg-Lee, K. Peterson, and D. Sharma for helpful comments on the manuscript.

## EXTENDED DATA

**Extended Data Figure 1:**
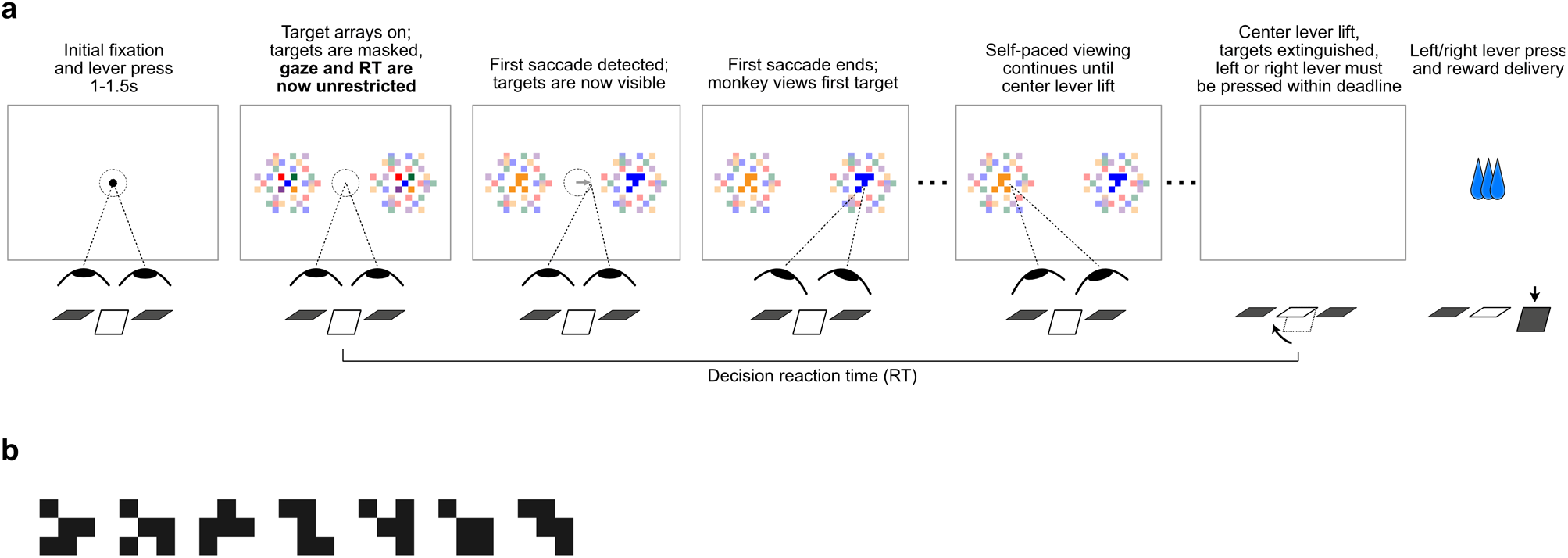
Task sequence and target shape primitives. **(a)** Illustration of the complete task sequence. Note that for illustration purposes the background is shown as white, and the surrounding crowders are shown at reduced saturation relative to the centrally located targets. However, on the actual task display, the background was dark, and target luminance was reduced relative to the crowders (see Fig. 1c), in order to better obscure the targets when they are not being viewed directly. **(b)** The 7 shape primitives used to generate choice stimuli. To create a new stimulus set, four shapes are selected without replacement from the 7 primitives; then each is rotated randomly by 0, 90, 180, 270 degrees. The four rotated shapes are then combined with four colors selected randomly from equally-spaced points on a color wheel defined within the RGB gamut in the CIELUV color space. A sub-set of 12 targets is then selected, and assigned to reward values ranging from 1-5 drops (Fig. 1b).

**Extended Data Figure 2:**
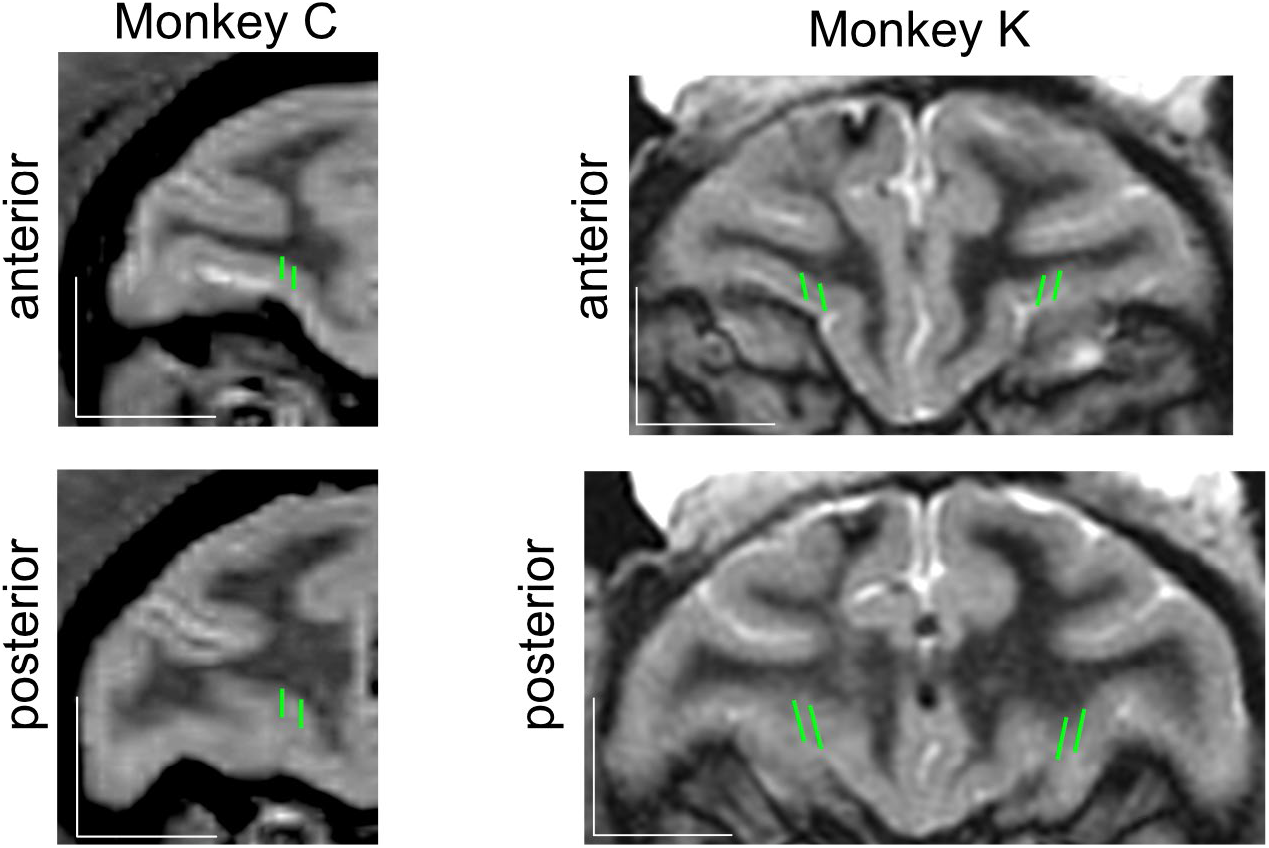
Recording locations. OFC recordings targeted the medial orbital gyrus, as well as the lateral bank of the medial orbital sulcus, over an anterior-posterior range of approximately +32 to +38mm relative to the intra-aural landmark. A representative anterior and posterior MR section (T2-weighted) is shown for each animal, with representative array trajectories shown in green. In Monkey C, recordings were from the left hemisphere only; in Monkey K, recordings were typically obtained bi-laterally. Scale bars are 10mm.

**Extended Data Figure 3:**
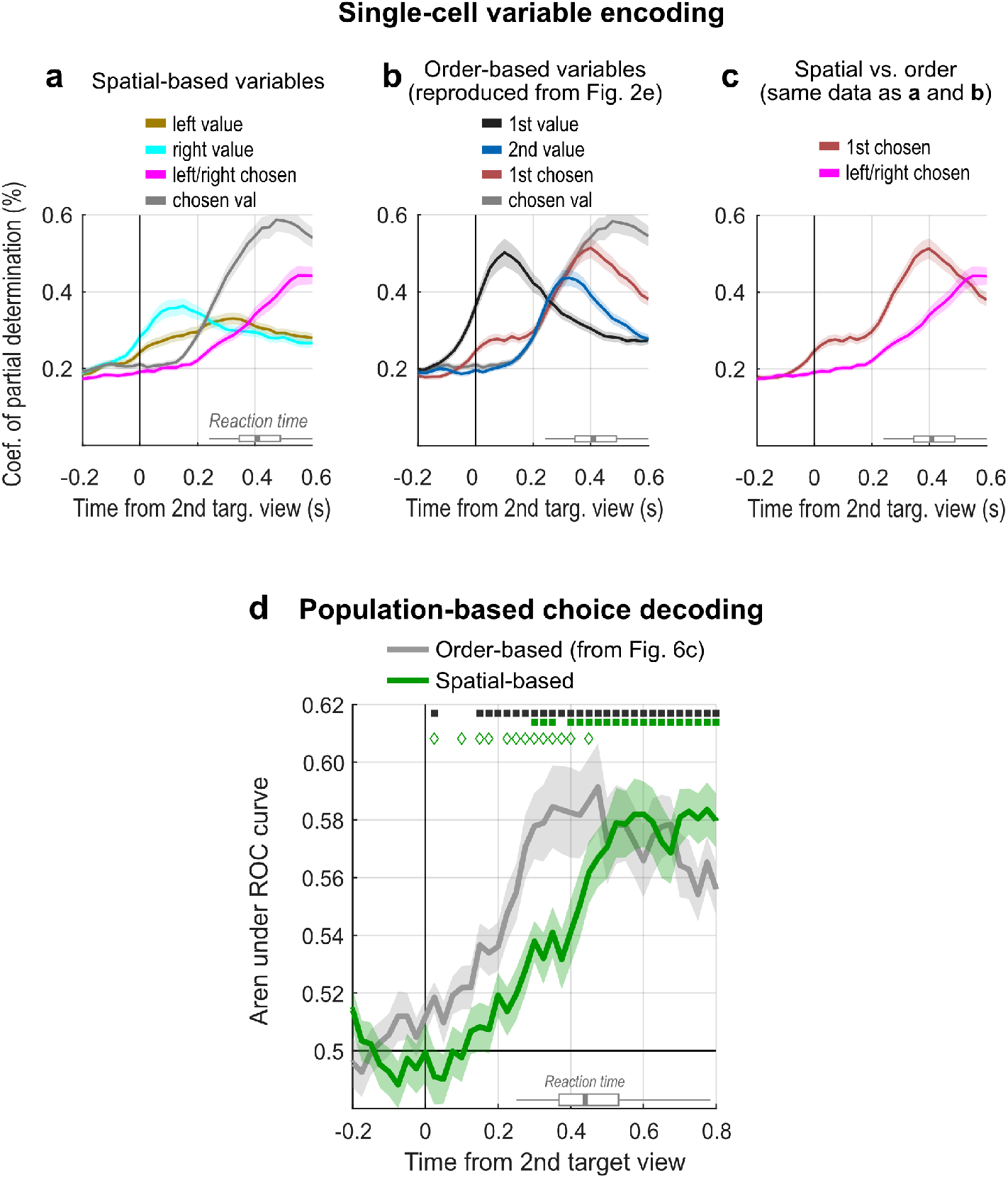
Order vs. spatial reference frame: **(a)** The encoding of spatial variables by single cells was measured using the CPD, as in Fig. 2d-e (see Methods). The variables were the value of the left offer (1-5 drops of juice), value of the right offer (1-5 drops), whether the left or right offer was chosen (Boolean), and the chosen value (1-5 drops). Means and SEMs are over 1450 single cells. **(b)** Data reproduced from Fig. 2e showing encoding of order-based variables in 1450 single cells. **(c)** Encoding of the variable ‘1st chosen’ from panel (b) plotted with the encoding of the variable ‘left/right chosen’ from panel (a). **(d)** Decoding of choice outcomes in a spatial reference frame (green) compared to order-based decoding (gray, data taken from Fig. 6c, “all dims”). Decoding was performed using the same approach as in Fig. 6c, except using spatial variables, i.e.: data were first residualized with respect to the identities of the left and right offer values, and held out trials were then decoded in terms of whether the left or right offer was chosen. Filled significance indicators: corrected p< 0.05 compared to 0.5; open significance indicators: uncorrected p< 0.05 comparing the gray and green lines. Otherwise, conventions are the same as in Fig. 6c.

**Extended Data Figure 4:**
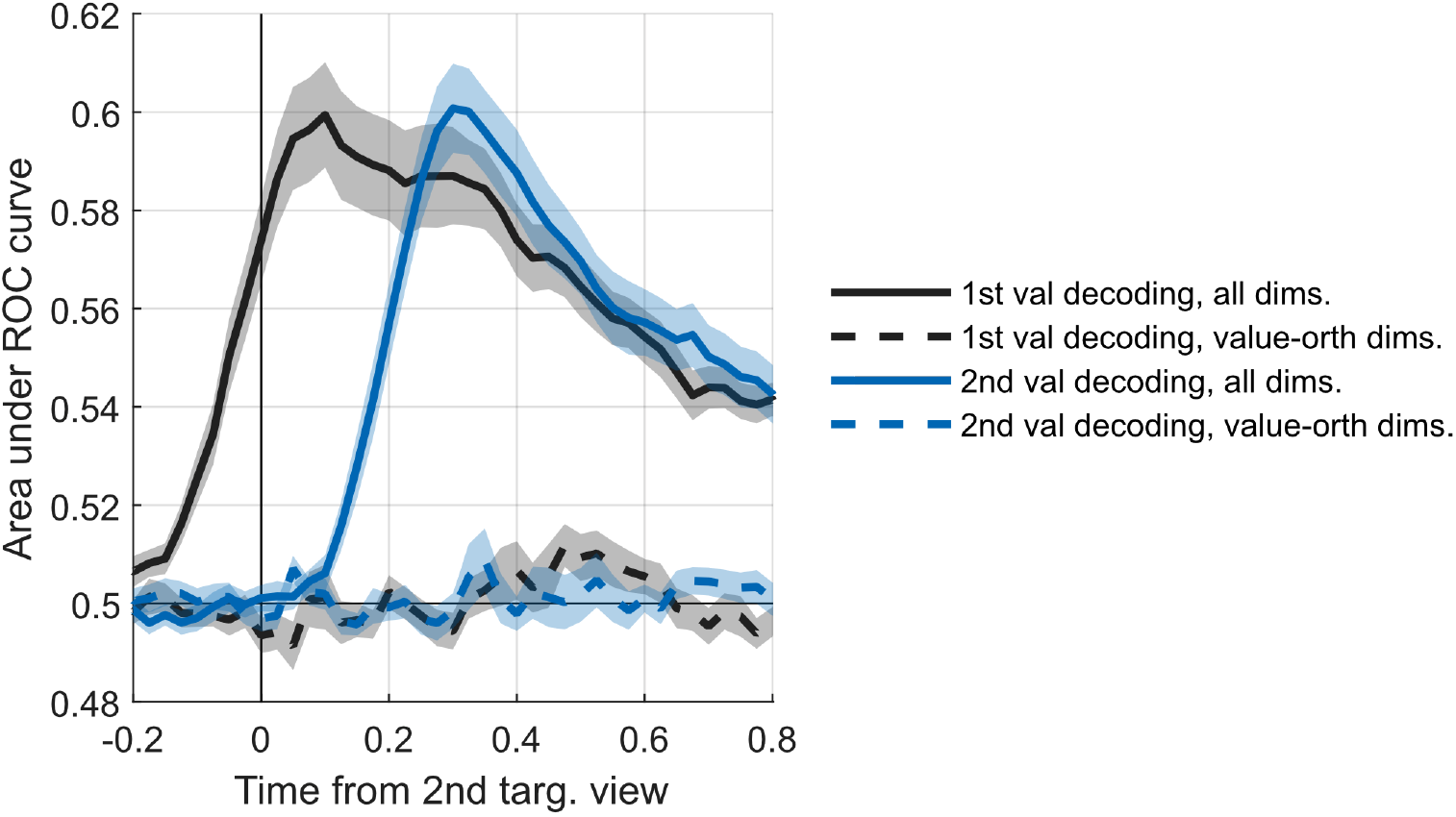
**Decoding of offer values in held out test trials**, using decoding models fit to ‘1st value’ and ‘2nd value’. Unlike the decoding of choice outcomes, decoding of values was performed using the non-normalized 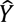 from Eqn. 3, rather than the condition-normalized NVEs. Within each session, classification was performed for adjacent levels of offer values (1 drop vs. 2 drops, 2 vs. 3, 3 vs. 4, and 4 vs. 5) and then averaged across the four adjacent-level classifications. The lines and shading indicate the mean and SEM of decoding accuracy across 32 sessions. The solid lines give the results of fitting, decoding, and classification performed over the native spiking data, utilizing all of the neural dimensions. The dashed lines give the results when fitting, decoding, and classifying data that were first projected into the subspace orthogonal to subspaces for ‘1st value’ and ‘2nd value’ (i.e. the subspace orthogonal to the plane in Fig. 6a, defined by weights obtained from Eqn. 2.) The fact that value cannot be decoded outside of the two main value dimensions indicates that the representation of value is low-dimensional. In comparison, significant decoding of choice outcomes *is* evident outside of the two main value dimensions (Fig. 6c, orange).

**Extended Data Figure 5:**
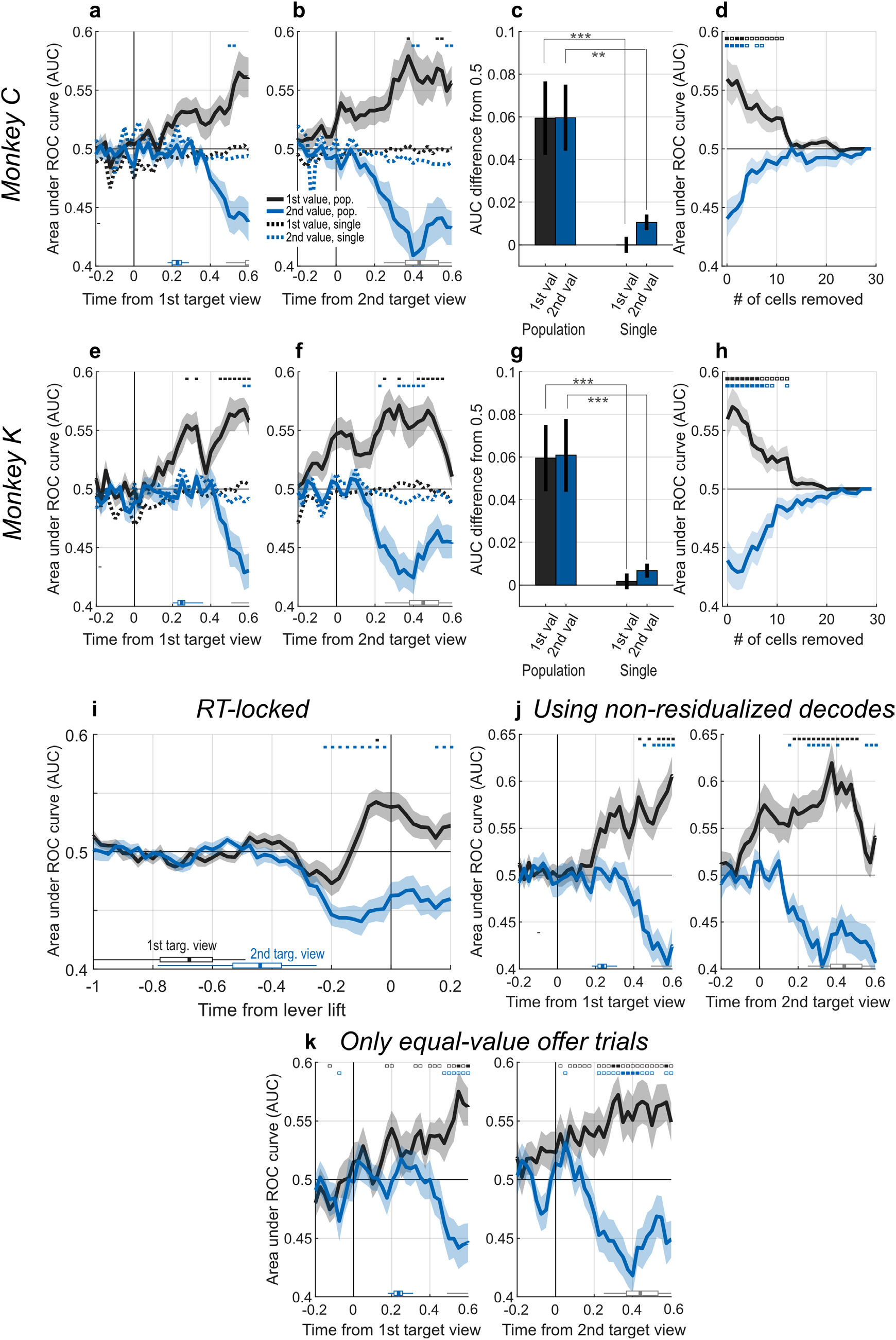
Results from individual monkeys and various controls. **(a-h)** Choice decoding and neuron dropping analyses for individual monkeys. Same format and conventions as Fig. 3. In panels (c) and (g), ** and *** indicate significant differences at uncorrected thresholds of p < 0.01 and p < 0.001, respectively. **(i)** The same data used to generate Fig. 3a-b, except time-locked to the decision RT, defined as the lift of the center lever. Note that when the data are aligned to the RT the first and second target viewing times (black and blue boxplots, respectively) are widely spread in time and have significant overlap. This overlap creates an artifact in which the area under the ROC curve for the ‘1st value’ decoder briefly goes below 0.5 (whereas in Fig. 3a-b it is always above 0.5). This is because within this brief time window, the encoding subspaces for ‘1st value’ and ‘2nd value’ are aligned (not shown), such that neural activity related to ‘2nd value’ projects onto the ‘1st value’ subspace. In other words, the ‘1st value’ decoder becomes contaminated by spiking activity related to the second offer value, resulting in predictions driven by the value of the second offer (i.e. that are below 0.5). Note that the subspace alignment and resulting contamination arises only because of the large spread between the first and second target viewing times when the data are RT locked. When the data are locked onto target viewing, the decoder weights for ‘1st value’ and ‘2nd value’ are uncorrelated (Fig. 4a), indicating orthogonal decoding subspaces **(j)** Choice decoding and neuron dropping analyses when using non-condition-normalized data to calculate the area under the ROC curve^36^. In this method, trials were grouped according to 66 unique conditions, defined as the combination of the first and second target identities. The non-condition-normalized data (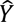 from Eqn. 3) were then used to calculate a separate AUC within each condition. Finally, within each session the AUCs were averaged across conditions to compute the session-wise AUC. AUCs could not be calculated for conditions without at least one trial of each choice outcome (first or second offer chosen); because there were 66 unique conditions, there were often too few trials per condition, and as a result ∼65% of test trials were discarded. For this reason, the main analysis uses normalized decodes, so that data from all eligible test trials can contribute to the AUC calculation. All conventions are the same as in Fig. 3a-b. **(k)** : Choice decoding as in Fig. 3a-b was performed using only those test trials in which the offers were equal in value (mean 43.9 trials in 16 sessions for Monkey K and 51.7 trials in 16 sessions for Monkey C). Filled significance indicators: corrected p<0.05 compared to 0.5; open indicators: uncorrected p<0.05 compared to 0.5. Otherwise, conventions are the same as in Fig. 3a-b

**Extended Data Figure 6:**
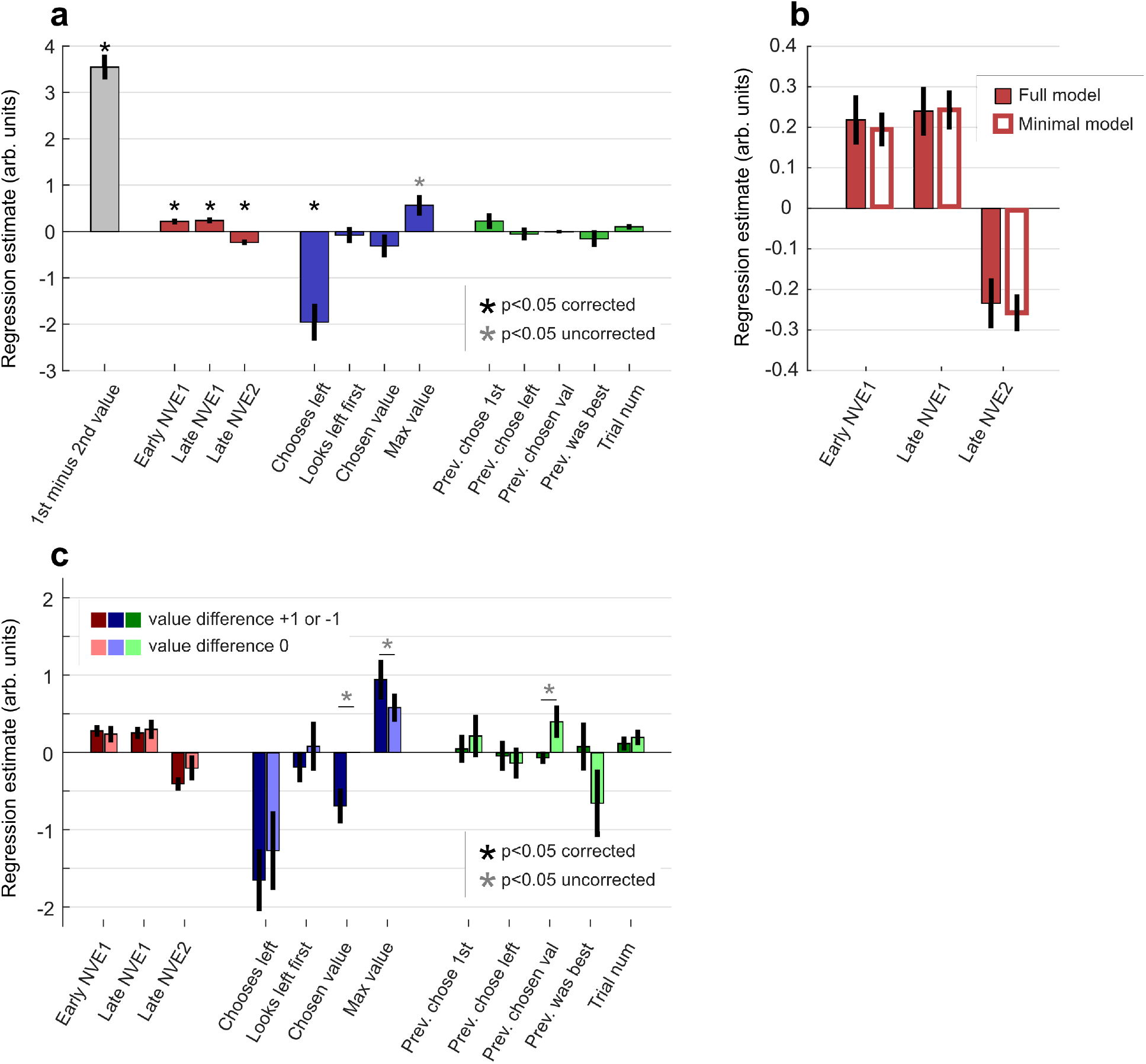
Neural-behavioral logistic model: Logistic regression was used to explain the choice of first vs. second offer with neural and behavioral variables described in Eqn. 4. **(a)** Mean and SEM of regression estimates over 32 sessions. Estimates are in arbitrary units. Significance indicators indicate difference from zero by t-test. **(b)** Estimates from neural signal variables in panel (a) (full model) were no different from estimates from a model containing only the first four terms in Eqn. 4 (minimal model). **(c)** Estimates from models fit using only trials with value difference of 1 drop compared to estimates fit using only trials with equal-value offers (dark vs. light bars). Significance indicators give difference between dark and light bar in each pair by paired t-test. No significant differences were found after p-value correction for multiple comparisons.

**Extended Data Figure 7:**
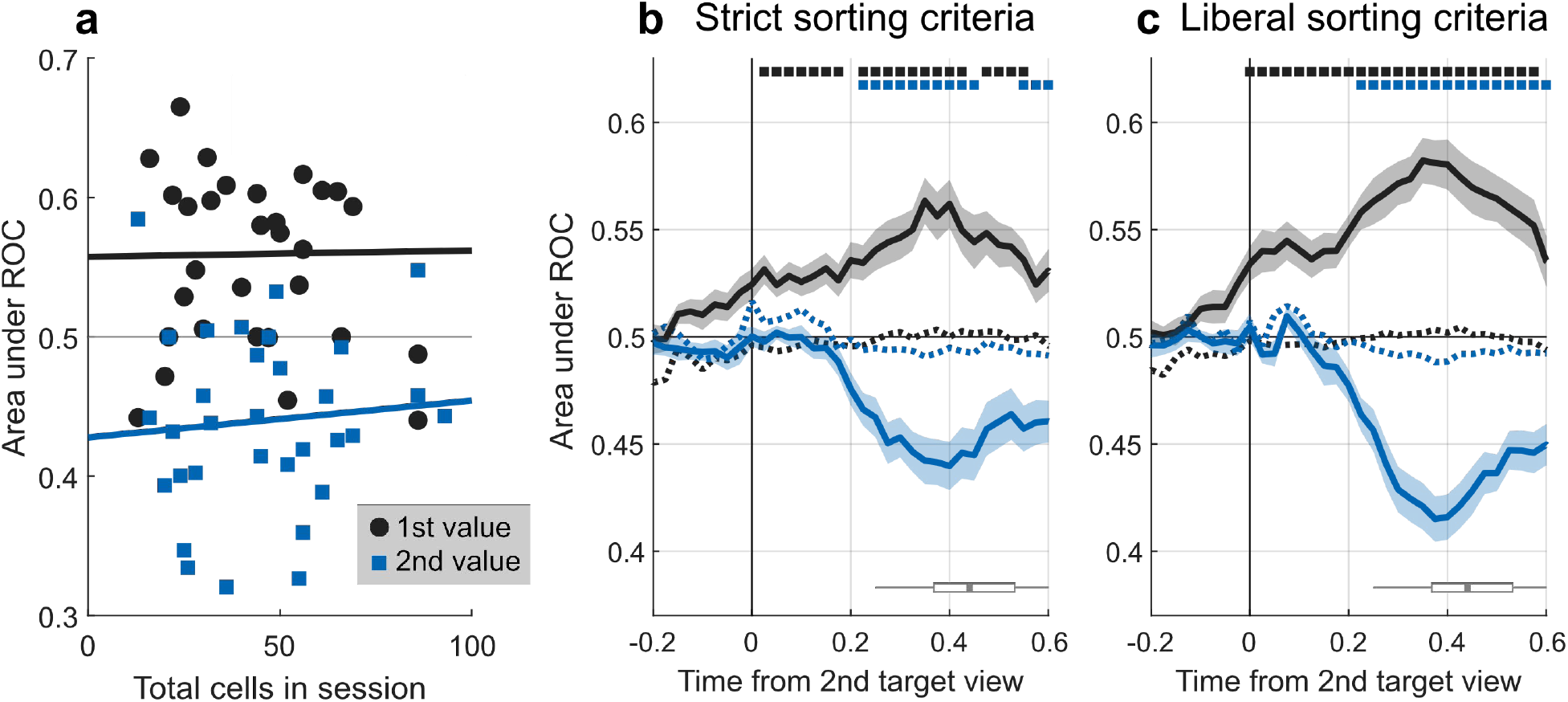
Decoding performance and number of cells used. **(a)** Choice decoding accuracy as a function of the number of cells recorded in each session, using the data from the ‘late’ epoch in Fig. 3b. Solid lines give linear fit. For ‘1st value’ the correlation was 0.015 (p = 0.94), and for ‘2nd value’ was 0.087 (p = 0.64). **(b-c)** Choice decoding effects referenced to second target viewing, using cells meeting ‘strict’ sort quality criteria (<1% of ISIs shorter than 2ms, and <10% of ISIs estimated to come from rogue spikes) or ‘liberal’ sorting criteria (<50% of ISIs from rogue spikes, no restriction on ISIs shorter than 2ms). Mean cells per session for strict criteria were 23.9, and for liberal criteria were 77.6. Conventions are the same as in Fig. 3b.

**Extended Data Figure 8:**
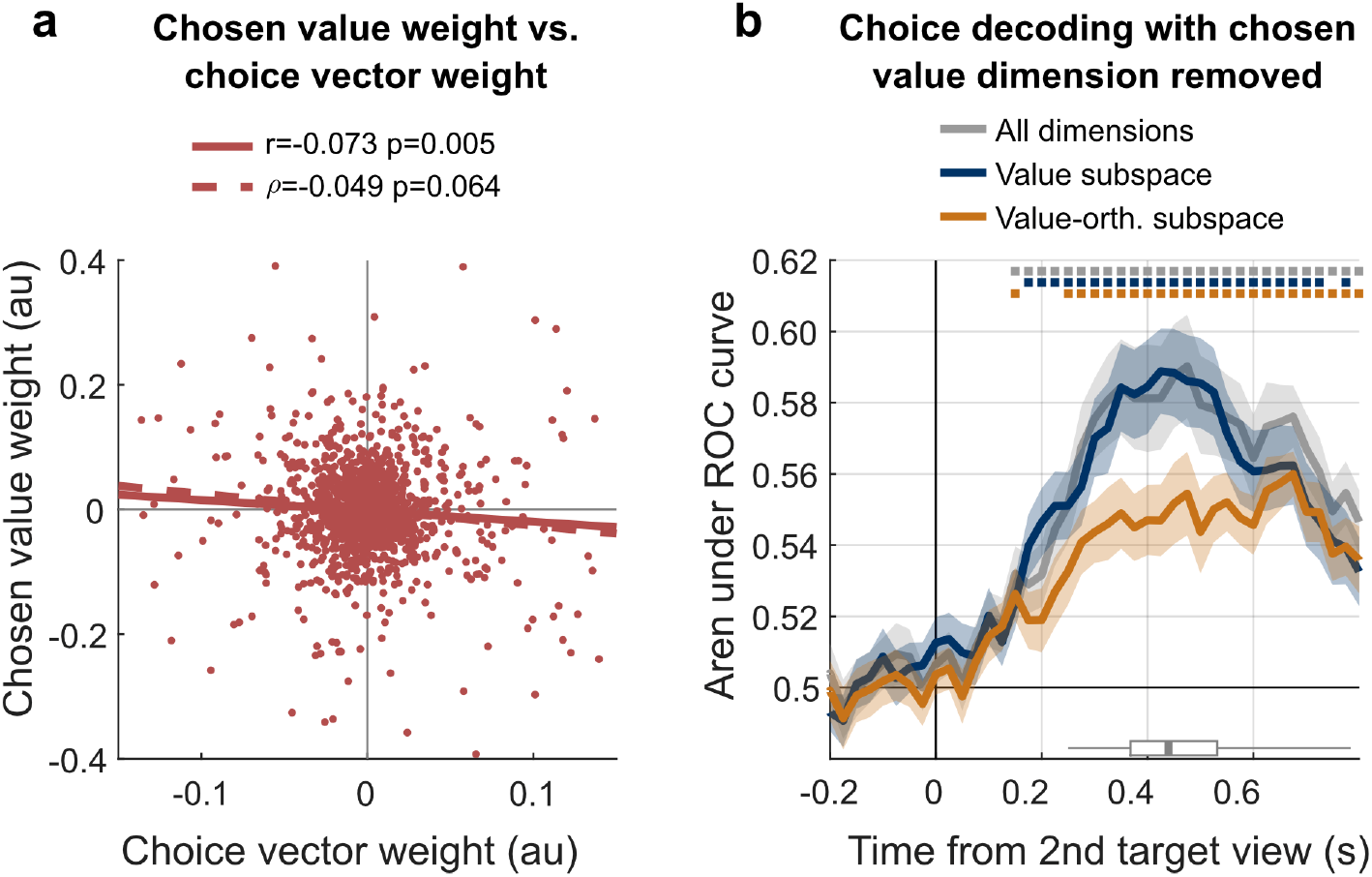
Chosen value relationship with choice decoding. **(a)** Data are taken from the late epoch in Fig. 3b. Solid lines indicate ordinary least squares fit; dashed lines indicate robust fit from MATLAB function `robustfit` with default arguments. Statistics *r* and *ρ* indicate Pearson’s and Spearman’s coefficients, respectively. The x-axis gives the per-cell weights that define the choice subspace (see Methods), and the y-axis gives the per-cell weights obtained by fitting Eqn. 2 to the variable ‘*chosen value*’ – i.e. the chosen value weights. **(b)** The choice decoding analysis in Fig. 6c was repeated after first projecting the spiking data into a subspace that is orthogonal to the subspace encoding chosen values. This was done with the same procedure used to project data into the subspace orthogonal to offer values, except that the matrix ***B*** has only one column, given by weights obtained by fitting Eqn. 2 to the variable ‘*chosen value*’ (the same weights as in panel **a**). This effectively removed information about chosen values from the spiking data, after which the other steps necessary to reproduce Fig. 6c were performed. Conventions are as in Fig. 6c.

**Extended Data Figure 9:**
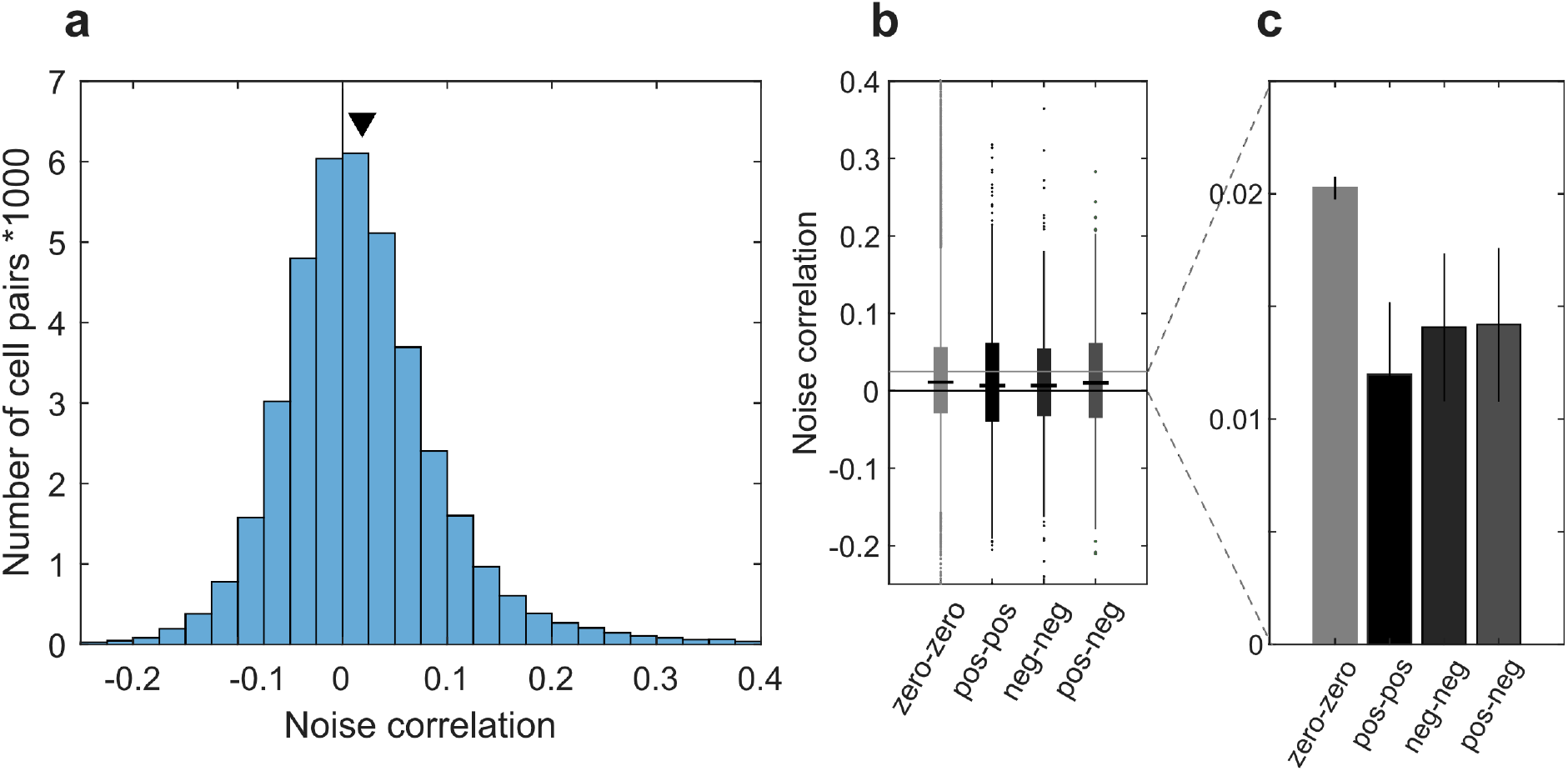
Pairwise noise correlations. measured from spike counts over a 200ms window centered 200ms after fixation onto the first target. This window was chosen because it contains the bulk of the response to the first-viewed target, but entirely excludes encoding of the second viewed target (see Fig. 2d). Similar results were obtained when using a 500ms window centered 250ms before targets onset (not shown). Before calculation of the noise correlations, spike counts were first z-scored according to the identity of the first target and whether the first fixation location was left or right. Cells weights for first offer value were obtained by Eqn. 2 with LASSO regularization, and designated as zero, positive, or negative according to the sign of the weights. **(a)** Histogram of noise correlations for all simultaneously recorded cells pairs (n=39,049). **(b)** Box plots of noise correlations between pairs of cells defined by the sign of the encoding weight for ‘1st value’. N = 26,734 cell pairs for zero-zero; n=718 for pos-pos; n=589 for neg-neg; n=537 for pos-neg. The horizontal line at 0.25 indicates the vertical range of the bar plot in panel C. **(c)** Same data as in (b) plotted as means and standard errors. A one-way ANOVA indicated a significant effect of group (F_3,28536_ = 4.2, p = 0.006).

## REFERENCES

1. Britten, K. H., Newsome, W. T., Shadlen, M. N., Celebrini, S. & Movshon, J. A. A relationship between behavioral choice and the visual responses of neurons in macaque MT. Vis. Neurosci. 13, 87–100 (1996).

2. Salzman, C. D., Britten, K. H. & Newsome, W. T. Cortical microstimulation influences perceptual judgements of motion direction. Nature 346, 174–177 (1990).

3. Yates, J. L., Katz, L. N., Levi, A. J., Pillow, J. W. & Huk, A. C. A simple linear readout of MT supports motion direction-discrimination performance. J. Neurophysiol. 123, 682–694 (2019).

4. Katz, L. N., Yates, J. L., Pillow, J. W. & Huk, A. C. Dissociated functional significance of decision-related activity in the primate dorsal stream. Nature 535, 285–288 (2016).

5. Gold, J. I. & Shadlen, M. N. The Neural Basis of Decision Making. Annu. Rev. Neurosci. 30, 535–574 (2007).

6. Shadlen, M. N. & Newsome, W. T. Neural Basis of a Perceptual Decision in the Parietal Cortex (Area LIP) of the Rhesus Monkey. J. Neurophysiol. 86, 1916–1936 (2001).

7. Cohen, M. R. & Newsome, W. T. Estimates of the Contribution of Single Neurons to Perception Depend on Timescale and Noise Correlation. J. Neurosci. 29, 6635–6648 (2009).

8. Ruff, D. A., Ni, A. M. & Cohen, M. R. Cognition as a Window into Neuronal Population Space. Annu. Rev. Neurosci. 41, 77–97 (2018).

9. Thorpe, S. J., Rolls, D. E. T. & Maddison, S. The orbitofrontal cortex: Neuronal activity in the behaving monkey. Exp. Brain Res. 49, 93–115 (1983).

10. Tremblay, L. & Schultz, W. Relative reward preference in primate orbitofrontal cortex. Nature 398, 704–708 (1999).

11. Padoa-Schioppa, C. & Assad, J. A. Neurons in the orbitofrontal cortex encode economic value. Nature 441, 223–226 (2006).

12. Rich, E. L. & Wallis, J. D. Decoding subjective decisions from orbitofrontal cortex. Nat. Neurosci. advance online publication, (2016).

13. Yamada, H., Louie, K., Tymula, A. & Glimcher, P. W. Free choice shapes normalized value signals in medial orbitofrontal cortex. Nat. Commun. 9, 162 (2018).

14. Ballesta, S., Shi, W., Conen, K. E. & Padoa-Schioppa, C. Values Encoded in Orbitofrontal Cortex Are Causally Related to Economic Choices. http://biorxiv.org/lookup/doi/10.1101/2020.03.10.984021(2020) doi:10.1101/2020.03.10.984021.

15. Knudsen, E. B. & Wallis, J. D. Closed-Loop Theta Stimulation in the Orbitofrontal Cortex Prevents Reward-Based Learning. Neuron (2020) doi:10.1016/j.neuron.2020.02.003.

16. Murray, E. A., Moylan, E. J., Saleem, K. S., Basile, B. M. & Turchi, J. Specialized areas for value updating and goal selection in the primate orbitofrontal cortex. eLife 4, e11695 (2015).

17. Rudebeck, P. H., Saunders, R. C., Prescott, A. T., Chau, L. S. & Murray, E. A. Prefrontal mechanisms of behavioral flexibility, emotion regulation and value updating. Nat. Neurosci. 16, 1140–1145 (2013).

18. Setogawa, T. et al. Neurons in the monkey orbitofrontal cortex mediate reward value computation and decision-making. *Commun*. Biol. 2, 126 (2019).

19. Eldridge, M. A. G. et al. Disruption of relative reward value by reversible disconnection of orbitofrontal and rhinal cortex using DREADDs in rhesus monkeys. Nat. Neurosci. 19, 37– 39 (2016).

20. Kennerley, S. W., Dahmubed, A. F., Lara, A. H. & Wallis, J. D. Neurons in the Frontal Lobe Encode the Value of Multiple Decision Variables. J. Cogn. Neurosci. 21, 1162–1178 (2008).

21. McGinty, V. B., Rangel, A. & Newsome, W. T. Orbitofrontal Cortex Value Signals Depend on Fixation Location during Free Viewing. Neuron 90, 1299–1311 (2016).

22. Hunt, L. T. et al. Triple dissociation of attention and decision computations across prefrontal cortex. Nat. Neurosci. 21, 1471–1481 (2018).

23. Padoa-Schioppa, C. Neuronal Origins of Choice Variability in Economic Decisions. Neuron 80, 1322–1336 (2013).

24. Kimmel, D. L., Elsayed, G. F., Cunningham, J. P. & Newsome, W. T. Value and choice as separable and stable representations in orbitofrontal cortex. Nat. Commun. 11, 3466 (2020).

25. Conen, K. E. & Padoa-Schioppa, C. Neuronal variability in orbitofrontal cortex during economic decisions. J. Neurophysiol. 114, 1367–1381 (2015).

26. Haefner, R. M., Gerwinn, S., Macke, J. H. & Bethge, M. Inferring decoding strategies from choice probabilities in the presence of correlated variability. Nat. Neurosci. 16, 235–242 (2013).

27. Zohary, E., Shadlen, M. N. & Newsome, W. T. Correlated neuronal discharge rate and its implications for psychophysical performance. Nature 370, 140–143 (1994).

28. Averbeck, B. B., Latham, P. E. & Pouget, A. Neural correlations, population coding and computation. Nat. Rev. Neurosci. 7, 358–366 (2006).

29. Enel, P., Wallis, J. D. & Rich, E. L. Stable and dynamic representations of value in the prefrontal cortex. eLife 9, e54313 (2020).

30. Crowder, E. A. & Olson, C. R. Macaque monkeys experience visual crowding. J. Vis. 15, 14 (2015).

31. Whitney, D. & Levi, D. M. Visual crowding: a fundamental limit on conscious perception and object recognition. Trends Cogn. Sci. 15, 160–168 (2011).

32. Fetsch, C. R. The importance of task design and behavioral control for understanding the neural basis of cognitive functions. Curr. Opin. Neurobiol. 37, 16–22 (2016).

33. Strait, C. E., Blanchard, T. C. & Hayden, B. Y. Reward Value Comparison via Mutual Inhibition in Ventromedial Prefrontal Cortex. Neuron doi:10.1016/j.neuron.2014.04.032.

34. Rudebeck, P. H., Ripple, J. A., Mitz, A. R., Averbeck, B. B. & Murray, E. A. Amygdala contributions to stimulus–reward encoding in the macaque medial and orbital frontal cortex during learning. J. Neurosci. 0933–16 (2017) doi:10.1523/JNEUROSCI.0933-16.2017.

35. Tibshirani, R. Regression Shrinkage and Selection via the Lasso. J. R. Stat. Soc. Ser. B Methodol. 58, 267–288 (1996).

36. Kang, I. & Maunsell, J. H. R. Potential confounds in estimating trial-to-trial correlations between neuronal response and behavior using choice probabilities. J. Neurophysiol. 108, 3403–3415 (2012).

37. Nienborg, H., R. Cohen, M. & >Cumming, B. G. Decision-Related Activity in Sensory Neurons: Correlations Among Neurons and with Behavior. Annu. Rev. Neurosci. 35, 463– 483 (2012).

38. Ruff, D. A. & Cohen, M. R. Simultaneous multi-area recordings suggest that attention improves performance by reshaping stimulus representations. Nat. Neurosci. 22, 1669– 1676 (2019).

39. Ni, A. M., Ruff, D. A., Alberts, J. J., Symmonds, J. & Cohen, M. R. Learning and attention reveal a general relationship between population activity and behavior. Science 359, 463– 465 (2018).

40. Moreno-Bote, R. et al. Information-limiting correlations. Nat. Neurosci. 17, 1410–1417 (2014).

41. Kohn, A., Coen-Cagli, R., Kanitscheider, I. & Pouget, A. Correlations and Neuronal Population Information. Annu. Rev. Neurosci. 39, 237–256 (2016).

42. Semedo, J. D., Zandvakili, A., Machens, C. K., Yu, B. M. & Kohn, A. Cortical Areas Interact through a Communication Subspace. Neuron 102, 249–259.e4 (2019).

43. Kaufman, M. T., Churchland, M. M., Ryu, S. I. & Shenoy, K. V. Cortical activity in the null space: permitting preparation without movement. Nat. Neurosci. 17, 440–448 (2014).

44. Jazayeri, M. & Afraz, A. Navigating the Neural Space in Search of the Neural Code. Neuron 93, 1003–1014 (2017).

45. Jurewicz, K., Sleezer, B. J., Mehta, P. S., Hayden, B. Y. & Ebitz, R. B. Irrational choices via a curvilinear representational geometry for value. 2022.03.31.486635 Preprint at https://doi.org/10.1101/2022.03.31.486635 (2022).

46. Yoo, S. B. M. & Hayden, B. Y. The Transition from Evaluation to Selection Involves Neural Subspace Reorganization in Core Reward Regions. Neuron 0, (2019).

47. Yoo, S. B. M., Sleezer, B. J. & Hayden, B. Y. Robust Encoding of Spatial Information in Orbitofrontal Cortex and Striatum. J. Cogn. Neurosci. 30, 898–913 (2018).

48. Wilson, R. C., Takahashi, Y. K., Schoenbaum, G. & Niv, Y. Orbitofrontal Cortex as a Cognitive Map of Task Space. Neuron 81, 267–279 (2014).

49. McNamee, D., Rangel, A. & O’Doherty, J. P. Category-dependent and category-independent goal-value codes in human ventromedial prefrontal cortex. Nat. Neurosci. 16, 479–485 (2013).

50. Vaidya, A. R., Sefranek, M. & Fellows, L. K. Ventromedial Frontal Lobe Damage Alters how Specific Attributes are Weighed in Subjective Valuation. Cereb. Cortex 1–11 doi:10.1093/cercor/bhx246.

51. Saez, R. A., Saez, A., Paton, J. J., Lau, B. & Salzman, C. D. Distinct Roles for the Amygdala and Orbitofrontal Cortex in Representing the Relative Amount of Expected Reward. Neuron 95, 70–77.e3 (2017).

52. Öngür, D. & Price, J. L. The Organization of Networks within the Orbital and Medial Prefrontal Cortex of Rats, Monkeys and Humans. Cereb. Cortex 10, 206–219 (2000).

53. Hill, D. N., Mehta, S. B. & Kleinfeld, D. Quality Metrics to Accompany Spike Sorting of Extracellular Signals. J. Neurosci. 31, 8699–8705 (2011).

54. Krajbich, I., Armel, C. & Rangel, A. Visual fixations and the computation and comparison of value in simple choice. Nat Neurosci 13, 1292–1298 (2010).

55. Lupkin, S. M. & McGinty, V. B. Monkeys exhibit human-like gaze biases in economic decisions. 2022.02.24.481847 Preprint at https://doi.org/10.1101/2022.02.24.481847 (2022).

